# Three major dimensions of human brain cortical ageing in relation to cognitive decline across the 8^th^ decade of life

**DOI:** 10.1101/2020.01.19.911420

**Authors:** SR Cox, MA Harris, SJ Ritchie, CR Buchanan, MC Valdés Hernández, J Corley, AM Taylor, JW Madole, SE Harris, HC Whalley, AM McIntosh, TC Russ, ME Bastin, JM Wardlaw, IJ Deary, EM Tucker-Drob

**Affiliations:** Lothian Birth Cohorts group, The University of Edinburgh, UK; Department of Psychology, The University of Edinburgh, UK; Scottish Imaging Network, A Platform for Scientific Excellence (SINAPSE) Collaboration, Edinburgh, UK; Division of Psychiatry, The University of Edinburgh, UK; Social, Genetic and Developmental Psychiatry Centre, King’s College London, London, UK; Centre for Clinical Brain Sciences, The University of Edinburgh, UK; Department of Psychology, University of Texas, Austin, Texas, USA; Alzheimer Scotland Dementia Research Centre, The University of Edinburgh, UK; UK Dementia Research Institute at The University of Edinburgh, UK

## Abstract

Different brain regions can be grouped together, based on cross-sectional correlations among their cortical characteristics; this patterning has been used to make inferences about ageing processes. However, cross-sectional brain data conflates information on ageing with patterns that are present throughout life. We characterised brain cortical ageing across the 8^th^ decade of life in a longitudinal ageing cohort, at ages ~73, ~76, and ~79 years, with a total of 1,376 MRI scans. Volumetric *changes* among cortical regions of interest (ROIs) were more strongly correlated (average *r* = 0.805, SD = 0.252) than were *cross-sectional* volumes of the same ROIs (average *r* = 0.350, SD = 0.178). We identified a broad, cortex-wide, dimension of atrophy that explained 66% of the variance in longitudinal changes across the cortex. Our modelling also discovered more specific fronto-temporal and occipito-parietal dimensions, that were orthogonal to the general factor and together explained an additional 20% of the variance. The general factor was associated with declines in general cognitive ability (*r* = 0.431, *p* < 0.001) and in the domains of visuospatial ability (*r* = 0.415, *p* = 0.002), processing speed (*r* = 0.383, *p* < 0.001) and memory (*r* = 0.372, *p* < 0.001). Individual differences in brain cortical atrophy with ageing are manifest across three broad dimensions of the cerebral cortex, the most general of which is linked with cognitive declines across domains. Longitudinal approaches are invaluable for distinguishing lifelong patterns of brain-behaviour associations from patterns that are specific to aging.

## Introduction

Accurately characterising patterns of brain ageing, alongside their determinants and functional significance, is a major challenge for developmental neuroscience. The eighth decade of life is a time in which risk for cognitive decline and dementia begins to accelerate markedly^1^, alongside consequent increases in personal and societal burden, and poorer quality of life^2–4^. The general under-representation of participants aged over 70 years in life-course brain imaging studies has been a problem for this research, along with the fact that many of our inferences about the progression of ageing-related brain changes come from cross-sectional datasets. Whereas cross-sectional information can potentially be informative for ageing, it has been strongly criticised in some quarters since it—unlike analysis of longitudinal data—is unable adequately to approximate the dimensionality and time-dependent dynamics of ageing^5–7^. Here, we investigate individual differences in patterns of cortical ageing using longitudinal data in a large sample of generally-healthy community-dwelling adults who were brain scanned three times, from their early-to-late 70s.

Older age is accompanied by a general decline in overall cerebral volume, with corresponding ventricular enlargement and increasing aggregation of other features such as white matter hyperintensites, and alterations in white matter microstructural properties^8–10^. At the more fine-grained regional level, cortical ageing is not uniform. Age effects appear to be stronger for some regions than others, with those areas ontogenetically and phylogenetically latest to develop—those that are more strongly linked to more complex cognitive functions—being those most affected^11^. However, it has become increasingly apparent that univariate accounts of brain ageing (considering a single brain area in isolation) are suboptimal for accurately characterising patterns of brain ageing. More recent accounts of brain organisation have used cross-sectional data to identify clusters of regions with shared morphometric characteristics^12–16^. Whereas such accounts have the potential to capture the coordinated patterns of age-related atrophy in health and disease, they are predominantly based on cross-sectional data, which, as noted above, cannot fully reflect the dynamics of within-individual change^17^. As such, the network configuration described by regional (cross-sectional) structural correlations is not necessarily optimal for evaluating brain ageing and its correlates.

Discovering the patterning of longitudinal changes in brain structure, and how such changes relate to longitudinal ageing-related cognitive decline, is crucial to understanding the neurobiology of cognitive ageing as distinct from the neurobiology of lifelong levels of cognitive function. Put simply, the fact that aspects of brain regions’ structure *correlate* together when measured on one occasion does not mean that they necessarily *change* together over time. Using longitudinal data to uncover the degree to which individuals show distinct patterns of cortical atrophy is likely to be valuable in the stratification of ageing subtypes against current and future cognitive and health outcomes, other biomarkers, and their potentially distinct determinants.

We are aware of only one previous study of multivariate longitudinal changes in cortical structure, which was conducted in an age-heterogeneous sample of participants with amnestic mild cognitive impairment (aMCI; N = 317)^18^. That study identified five groupings, each of which comprised regions with strongly correlated atrophic profiles. These broadly described 1) posterior default mode, 2) prefrontal, 3) medial temporal, 4) ‘spared’ (sensorimotor and occipital), and 5) a diffuse global atrophic pattern. The authors concluded that this might reflect multiple patterns of coordinated neuronal degradation with potentially distinct biological substrates among their clinical sample^18^. As yet, it remains unclear whether the presence of cross-sectional correlations between brain regions bears any relation to their shared patterns of change over time among non-clinical, generally-healthy older adults, whose neurobiological and cognitive changes may occur during early, prodromal phases of cognitive decline, when prevention and intervention efforts may be most likely to succeed.

In the present study, we used longitudinal data collected on three occasions across the eighth decade of life to characterise the patterns of cortical ageing in a group of community-dwelling older adults. We first estimated the cross-sectional levels (baselines) and longitudinal changes for each cortical region. We then explored their factor structure to discover any clusters of regions that exhibited correlated changes over time. We next related the factors of brain cortical change to changes in major age-sensitive domains of cognitive capability: memory, visuo-spatial reasoning, and processing speed. We placed particular emphasis on documenting how factors underlying cortex-wide and more regionally-specific constellations of variation in volumetric atrophy related to longitudinal ageing-related cognitive declines.

## Methods

### Participants

Participants were drawn from the Lothian Birth Cohort 1936^19–21^, a longitudinal study of brain and cognitive ageing in healthy community-dwelling older adults. The participants were initially recruited into this same-year-of-birth project at around 70 years of age (Wave 1, N = 1,091) where they underwent a number of cognitive, health and medical assessments. They were subsequently followed-up at ages ~73 (Wave 2, N = 866), ~76 (Wave 3, N = 697), and ~79 (Wave 4, N = 550), where they completed mostly the same tests as previously, with the addition of structural brain imaging assessments (see next section). Here, we included participants for Waves 2, 3 and 4, for whom concomitant brain imaging and cognitive data were available. Whole blood was drawn at baseline from which genomic DNA was isolated (at the Wellcome Trust Clinical Research Facility Genetics Core, Western General Hospital, Edinburgh). Participants provided written informed consent prior to testing at each wave. The LBC1936 study was approved by the Multi-Centre for Scotland (MREC/01/0/56), Lothian (LREC/2003/2/29) and Scotland A (07/MRE00/58) Research Ethics Committees.

### MRI Acquisition & Analysis

Brain structural MRI data were acquired according to a previously published protocol^22^ at Waves 2, 3 and 4. A 1.5T GE Signa HDxt clinical scanner (General Electric, Milwaukee, WI, USA) with a manufacturer-supplied eight-channel phased-array head coil was used to acquire 3D T1-weighted volumes in the coronal orientation at 1mm isotropic resolution. All scans were assessed by a consultant neuroradiologist, which included assessment of evidence for stroke. Acquired volumes were then processed in FreeSurfer v5.1^23–25^. This involved segmentation of each volume, identifying brain tissue types, followed by parcellation of cortical grey matter into 34 regions per hemisphere, according to the Desikan-Killiany atlas^26^ (Supplementary Figure 1). Output for each image was visually assessed for segmentation and parcellation errors, which were then corrected manually. Segmentations with errors that could not be corrected were excluded. Using FreeSurfer’s longitudinal processing stream^27^, each participant’s data across all waves were then resampled together, in order to minimise any erroneous longitudinal variation in parcellation. Regional volumes for each participant at each wave were then derived from the output of the longitudinal stream. The following analyses are based on 1,376 MRI scans (Wave 2 N = 629; Wave 3 N = 428; Wave 4 N = 319).

### Cognitive Measurement

Participants underwent a detailed battery of standardised cognitive tests that assessed major domains relevant to ageing. These were categorised into three cognitive domains based upon well-fitting, hierarchical structural equation models tested in our previously-published work^28,29^. Visuospatial ability was indicated by performance on Matrix Reasoning and Block Design from the Wechsler Adult Intelligence Scale III^UK^ (WAIS III^UK^)^30^, and the sum of Spatial Span Forward and Backward from the Wechsler Memory Scale III^UK^ (WMS III^UK^)^31^. Processing Speed was measured with Symbol Search and Digit Symbol Substitution from the WAIS III^UK^, Visual Inspection Time^32^, and Four-Choice Reaction Time^33^. Verbal Memory ability was ascertained using Logical Memory (sum of immediate and delayed) and Verbal Paired Associates (sum of immediate and delayed) from the WMS III^UK^, and Digit Span Backwards from the WAIS III^UK^. The Mini-Mental State Examination (MMSE)^34^ was also administered.

### APOE Genotyping

TaqMan technology was used to identify *APOE* e4 carriers. Status was determined according to genotyping on the two polymorphic sites (rs7412 and rs429358) that account for e2, e3 and e4 alleles^35^.

### Statistical Analysis

Figure 1 illustrates the analytical framework. First, we characterised the trajectories of cortical ageing at the level of each the 34 brain regions of interest (ROI). For each ROI per hemisphere, we fitted a separate growth curve in a structural equation modelling (SEM) framework in R^36^ using the ‘lavaan’ package^37^. We used full information maximum likelihood estimation throughout. The unstandardized estimate of slope was reported as % change per annum.

**Figure 1.**
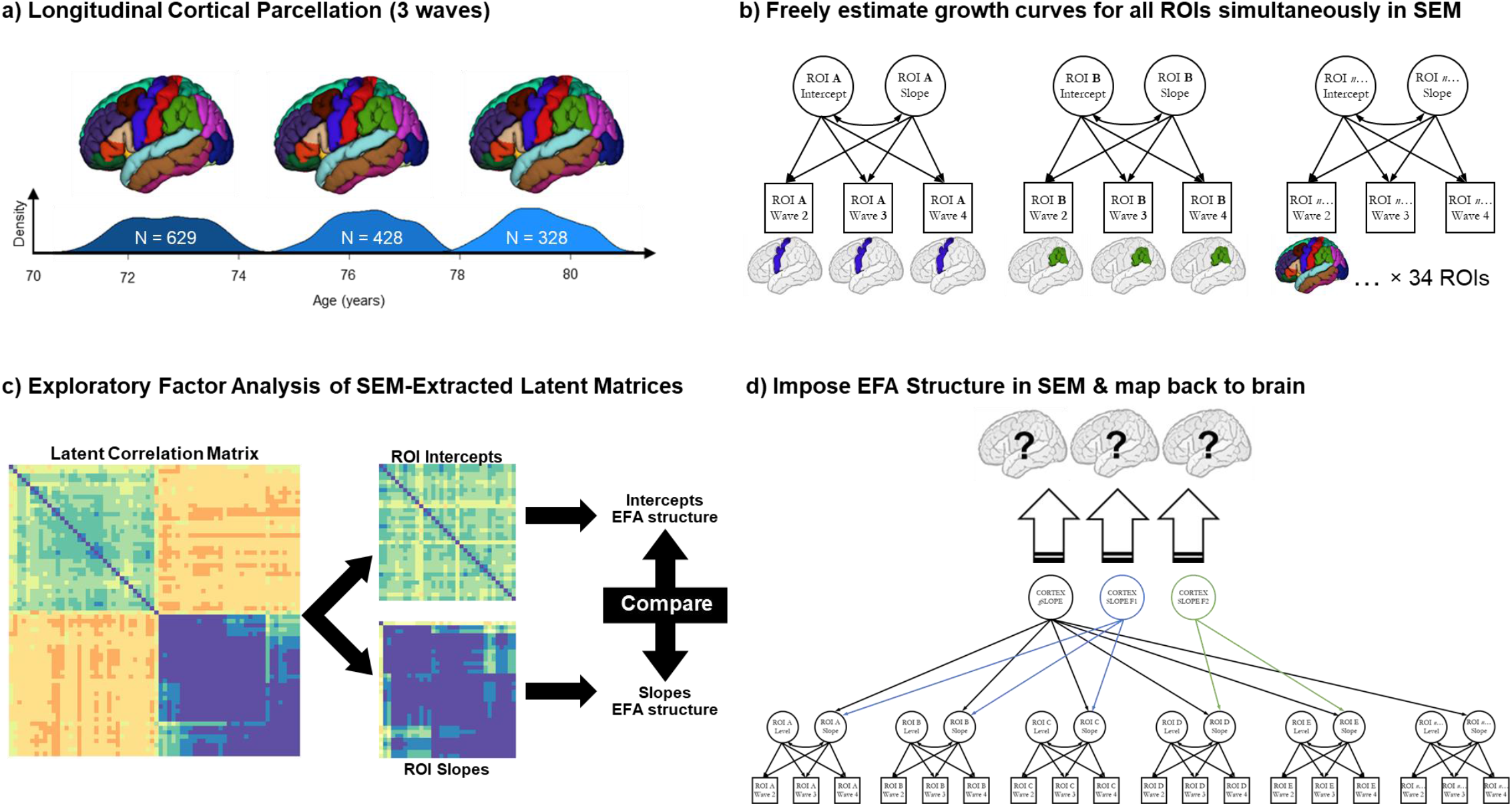
Analysis pipeline for establishing dimensions of brain cortical ageing. *Note.* **a)** T1-W brain MRI volumes were parcellated into 34 regions per hemisphere using the FreeSurfer longitudinal pipeline, at 3 waves from c.70-80 years old; **b)** we then simultaneously estimated growth curves (freely-estimated latent intercepts and slopes) for each region of interest (ROI) simultaneously, with a structural equation model (SEM); **c)** we extracted the resultant latent correlation matrix from the SEM in (b), separated these into intercepts and slopes used these matrices to investigate their correlational structure, using a Schmid-Leiman exploratory factor analyses (EFA). We ran formal tests to compare this structure across hemispheres (left v right) and between intercepts and slopes; **d)** we then conducted a confirmatory factor analysis (CFA); here we took the same model as in (b), but now imposed the three-factor structure implied by the EFA in (c), rather than freely estimating the slopes of each ROI. ROI latent intercepts were allowed to covary with all latent factors (not shown). The magnitude of the loadings for each of the factors was then mapped back onto the brain ROIs, indicating the groups of regions where atrophy is correlated, and allowing us to ask how these changes are correlated with genetic and cognitive status (e.g. Figure 4).

To investigate the possibility that there are spatially distinct dimensions of cortical ageing, our first step was to run an exploratory factor analysis (EFA) on the intercepts and slopes of the left hemisphere brain cortical ROIs (to be subsequently tested against the right hemisphere to test whether both hemispheres show similar patterns). First, we fitted a SEM in which growth curves (intercepts and slopes) for all 34 left hemisphere ROIs were freely estimated. We then extracted the estimated latent correlation matrix and separated it into two parts; one of intercepts and one of slopes. The ‘nearPD’ function (from the ‘psych’ package in R)^38^ was used to scale estimated values so that they were positive definite (ranging from −1 to 1, due to some residual variances initially having negative estimates).

Slope and intercept covariances were examined as hierarchically-clustered heatmaps. Based on this information, we conducted a Schmid-Leiman^39^ transformation (from the ‘psych’ package) to examine the oblique factor structure beyond any common variance shared (i.e. a bi-factor model). We used default parameters, comprising minimum residual OLS (fm = “minres”) and oblimin rotation (rotate = “oblimin”). This transformation was conducted separately for intercepts and slopes, retaining factor loadings >0.3. Given our primary interest was in slope covariances, we initially examined 2, 3 and 4 factor solutions for the slopes – selection of the optimal solution was based on eigenvalue magnitudes and distribution across factors, loading pattern (i.e. presence of factors with strong loadings were preferred), and the root mean square of residuals (RMSR). To test whether the identified correlation structure for left hemisphere intercepts and slopes was replicable (hemisphere differences in brain structural measures are extremely modest-to-null according to cross-sectional estimates^8,40,41^), we then repeated this exploratory analysis for the right hemisphere ROIs. We used Pearson’s *r* and the coefficient of factor congruence^42^ to formally quantify whether the resultant factor loading pattern replicated across hemispheres. We also ran these same two comparisons between the factor structure of intercepts and slopes within each hemisphere; that is, we tested whether brain cortical *level* correlations resembled cortical *change* correlations.

Although all participants were free from dementia diagnosis at baseline recruitment, we conducted a sensitivity analysis to ascertain whether the observed correlational structure may have been driven by participants who may have subsequently developed dementia or cognitive impairment. We did so by removing all those who had either, i) subsequently reported having received a diagnosis of dementia at any wave, or ii) had a score <24 on the MMSE at any wave. We then compared the resultant loading patterns with the results from the whole group analysis, using Pearson’s *r* and the coefficient of factor congruence, as above. We conducted a second sensitivity analysis, repeating these steps, this time removing participants whose MRI scans indicated stroke, as assessed by a consultant neuroradiologist (author JMW).

Next, we conducted a confirmatory factor analysis, imposing the slope factor structure identified from the Schmid-Leiman transform. That is, in a confirmatory model, we imposed the factor loading pattern identified from the the Schmid-Leiman-transformed EFA to estimate the loadings of each ROI’s slope factor on the relevant slope factor of cortical change. We specified the group factors to be uncorrelated with the general factor, but allowed the group factors to correlate with one another. Given that the EFA was conducted on correlation matrices rather than a data frame itself, this step was necessary to allow us to investigate correlates of the observed factors of change. Growth curve intercepts for each ROI were freely-estimated, and allowed to correlate with all other latent variables. Where the loadings of any ROI slope onto any slope factor were non-significant (*p* < 0.05), these were set to zero.

We then extended these multivariate SEMs to analyse the degree to which the factors of cortical change were associated with trajectories of cognitive change. We examined associations at the level of general cognitive ability (‘*g*’; see Supplementary Figure 2 and ^43^), and for the correlated cognitive domains of visuospatial, processing speed and memory, as well as with *APOE* e4 allele carrier status. The levels and changes in cognitive function were estimated in a Factor of Curves growth curve SEM (whereby each cognitive test over time has its own intercept and slope, and these contribute to an overall latent intercept and slope of all cognitive tests in that domain)^44^. We fitted one model for each of the cognitive analyses (*g*, visuospatial, processing speed, memory) in association with our model of cortical change. To aid model convergence and ensure construct consistency, we fixed factor loadings for both the cognitive and MRI sides of the SEM according to our initial measurement models / confirmatory factor analyses (CFAs). Specifically, for the brain cortical aspects of these models, we fixed the ROI slope factor loadings on the three overall slope factors. For each of the four cognitive models (three cognitive domains and overall general cognitive model), we fixed the factor loadings of the cognitive test latent intercept and latent slope factors on the cognitive domain intercept and slope factors. The cognitive test intercepts and slopes were corrected for sex; the CFA of cortical change indicated that there were no sex differences in any cortical factor. Residual correlations were included between the slopes of spatially contiguous ROIs^18^, and the residual variance of regional slope factors whose estimates were negative were constrained to zero to allow the model to converge on in-bounds estimates.

Associations across all cortical factors and all cognitive domains were corrected for multiple comparisons using the False Discovery Rate (FDR)^45^. All analyses were conducted in R 3.5.0 (“Joy in Playing”)^36^, except for CFA and bivariate growth curve models, which were conducted using MPlus 8.2^46^.

## Results

### Global and Regional Brain Cortical Ageing

Participant characteristics at each wave are shown in Table 1. Those who attended subsequent waves of neuroimaging tended to have modestly higher cognitive baseline scores across waves (and slightly lower cortical volume at Wave 3 only; Supplementary Tables 1 and 2). Overall and regional reductions in cortical volume are shown in Figure 2 and Supplementary Table 3. Trajectories for each region are also plotted in Supplementary Figure 3. Overall cortical volume showed a significant decline of 0.87% per annum (3,475mm^3^ of baseline volume). However, the rate of annual decline was not consistent across regions. Areas of overall greatest change included frontal and temporal poles (≥ 1.30% reduction per annum) parietal cortex, lateral occipital and lateral aspects of the frontal lobe. In contrast, the insula, cingulate and pre- and post-central areas showed much lower volumetric declines (≥ 0.67% reduction per annum).

**Table 1.** Participant characteristics.

|  | **Wave 2** |  | **Wave 3** |  | **Wave 4** |  |
| --- | --- | --- | --- | --- | --- | --- |
|  | **M (SD)** | **N** | **M(SD)** | **N** | **M(SD)** | **N** |
| **Age (years)** | 72.49 (0.71) | 866 | 76.24 (0.68) | 697 | 79.32 (0.62) | 550 |
| **Sex M:F** | 448:418 | 866 | 360:337 | 697 | 275:275 | 550 |
| **APOE e4 carriers (N:Y)** | 575:245 | 820 | 459:200 | 659 | 365:156 | 521 |
| **Cortical Volume (mm<sup>3</sup>)** | 398508 (396537) | 629 | 393294 (36251) | 428 | 382675 (35046) | 328 |
| **Matrix Reasoning** | 13.17 (4.96) | 863 | 13.04 (4.91) | 689 | 12.90 (5.03) | 535 |
| **Block Design** | 33.64 (10.08) | 864 | 32.18 (9.95) | 691 | 31.20 (9.63) | 535 |
| **Spatial Span Total** | 14.69 (2.76) | 861 | 14.61 (2.73) | 690 | 14.13 (2.72) | 536 |
| **Symbol Search** | 24.61 (6.18) | 862 | 24.60 (6.46) | 687 | 22.73 (6.63) | 528 |
| **Digit Symbol** | 56.40 (12.31) | 862 | 53.81 (12.93) | 685 | 51.24 (13.01) | 535 |
| **Inspection Time** | 111.22 (11.79) | 838 | 110.14 (12.55) | 654 | 106.96 (13.6) | 465 |
| **4-Choice Reaction Time** | 0.65 (0.09) | 865 | 0.68 (0.10) | 685 | 0.71 (0.11) | 543 |
| **Logical Memory** | 74.23 (17.89) | 864 | 74.58 (19.20) | 688 | 72.71 (20.39) | 542 |
| **Verbal Pairs** | 27.18 (9.46) | 843 | 26.41 (9.56) | 663 | 27.14 (9.55) | 497 |
| **Digit Span Backwards** | 7.81 (2.29) | 866 | 7.77 (2.37) | 695 | 7.56 (2.18) | 548 |

*Note.* Across Wave 2 to 4 (brain imaging data was not collected at Wave 1 of the study).

**Figure 2.**
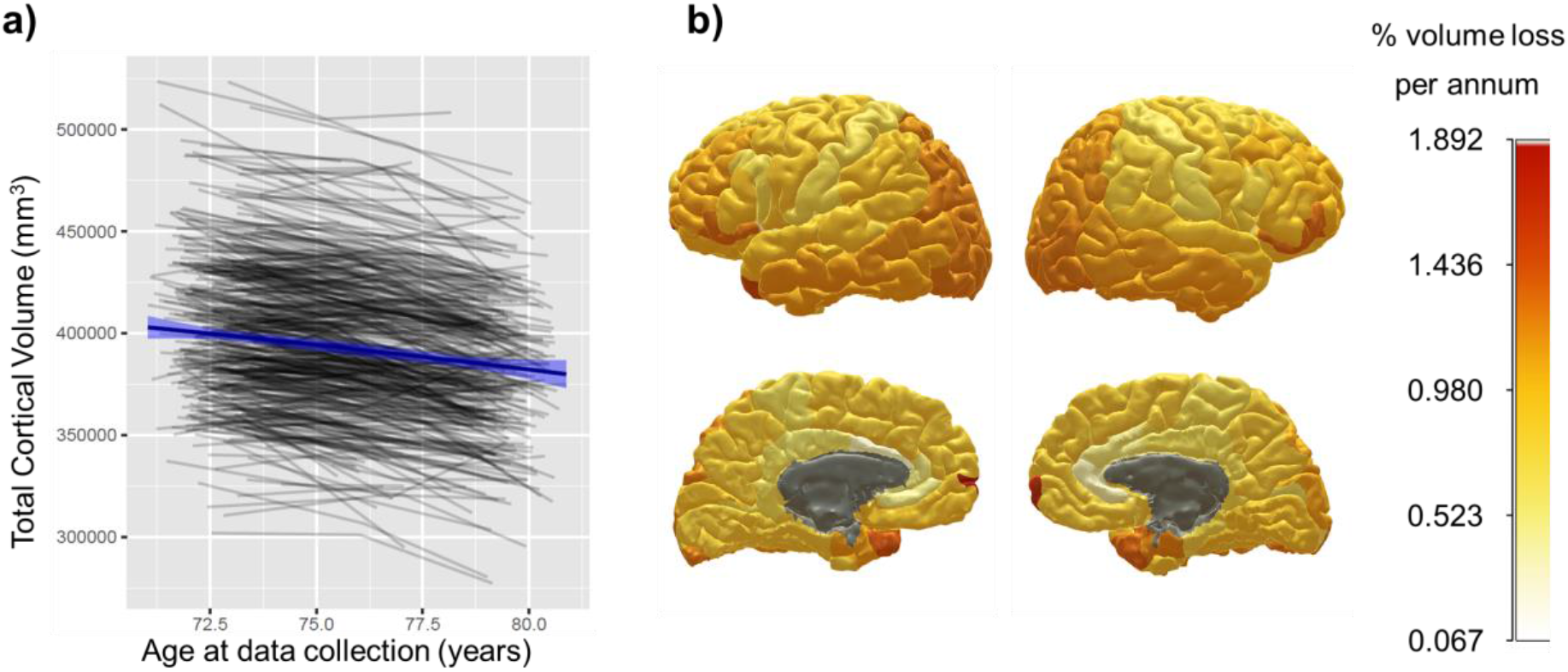
Global and regional cortical volumetric change from age 70 to 81 years of age. *Note.* **a)** shows, in grey lines, individual trajectories of global cortical volume; blue line denotes the mean linear trajectory with 95% CIs. **b)** shows the mean % loss per annum, estimated by growth curve models, for each cortical region of interest; warmer colours denote a steeper decline (grey areas indicate non-significant change). Individual plots by region are shown in Supplementary Figure 3.

### Factors of Brain Cortical Change

#### Exploratory Factor Analysis

Across the left hemisphere of the cortical mantle, the rates of the 34 ROIs’ cortical changes were highly correlated (average *r* = 0.81, SD = 0.25). These correlations of changes were higher than the correlations between the ROIs’ cross-sectional levels (average *r* = 0.35, SD = 0.18). Results of the exploratory Schmid-Leiman analyses are shown in Supplementary Figure 4, and in Supplementary Table 4. Consistent with the hierarchically-clustered heatmap, a solution for a general factor and two additional factors was selected; a three or four factor solution contained at least one factor with no loadings >0.3 and at least one factor with eigenvalues close to zero (Supplementary Tables 5 and 6; all RMSR < 0.05). Thirty-three of the 34 ROIs loaded on the first unrotated (general) factor, with only 4 of them < 0.5; the mean loading was 0.78. Beyond this general factor, ROI slopes showed two relatively distinct factors that were orthogonal to it; one had larger loadings in mainly fronto-temporal ROIs, and the other had high loadings in predominantly occipito-parietal ROIs. When we conducted the same Schmid-Leiman factor analysis on the right hemisphere data (Supplementary Table 7) and formally compared the factor structure between hemispheres, we found these two be highly similar (Supplementary Table 8; *r* range 0.61 to 0.78; factor congruence range 0.73 to 0.99). The factor structure for the ROIs’ intercepts was consistent between hemispheres. Importantly, however, the slope and intercept factor structure were less similar (*r* range 0.04 to 0.67; factor congruence range 0.45 to 0.93). Thus, the brain regions’ cross-sectional correlational structure showed relatively weak correspondence to the pattern of correlated changes across the cortex.

Given the high level of agreement between left and right hemispheres, we subsequently conducted a Schmid-Leiman analysis for bilateral regional averages (Figure 3 and Supplementary Table 9). A first (unrotated) general factor explained 55% of the variance among cortical volumetric slopes (loadings range 0.50 to 0.82, M = 0.75, SD = 0.09). The other two factors, which are independent of the general factor, accounted for 29% (loadings M = 0.58, SD = 0.13) and 13% (loadings M = 0.49, SD = 0.14) of the slope variance. In contrast, the same analysis of the cortical *intercepts* resulted in a factors that accounted for substantially less variance (General = 27%, Factor 1 = 16%, and Factor 2 = 3%). Again, the factor structure for intercepts and slopes was not similar; their lack of correspondence with the slope factor structure was confirmed through formal tests (Supplementary Table 10), indicating that although factor congruence of the General factor and Factor 1 were good (0.95 and0.88), this was not so for Factor 2 (0.54), nor were the factor loading correlations strong for any factor (*r=* 0.30, 0.53 and 0.54). Re-running these analyses once participants with a self-reported diagnosis of dementia or MMSE score > 24 (N = 31) or neuroradiologically-identified stroke (N = 119) at any wave had been removed did not substantially alter the patterns (Supplementary Tables 11 and 12, and Supplementary Figures 5 and 6).

**Figure 3.**
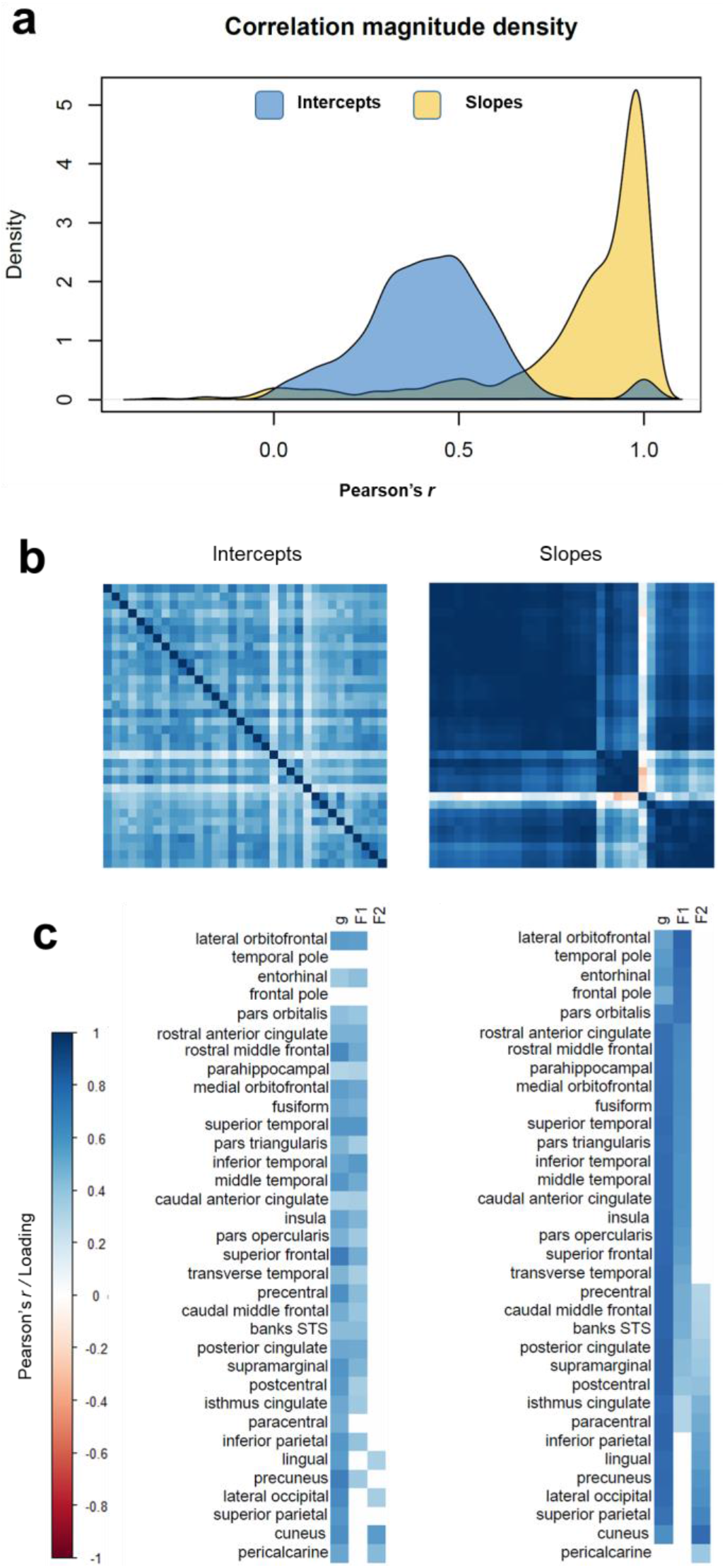
Correlation matrices and exploratory factor loadings for ROI intercepts and slopes, estimated using left-right average values for ROIs. *Note.* Exploratory factor analyses. **(a)** density plots of the correlation magnitudes among freely-estimated intercepts and slopes; **(b)** heatmaps of the correlations among freely-estimated latent intercepts (left) and slopes (right), intercept axes are fixed according to the hierarchically-clustered slope matrix; **(c)** loadings of each ROI’s intercept and slope on a general factor of cortical change (“*g*”) and on two additional factors identified from an exploratory Schmid-Leiman factor analysis, conducted on the same latent correlation matrices as shown in (b). Loadings reported in Supplementary Table 9. Factor F1 pertains to fronto-temporal, and F2 to occipito-parietal regions.

#### Confirmatory Factor Analysis – mapping back to the brain

We then undertook a confirmatory factor analysis (CFA) SEM. Here, we used the loading structure for the ROI changes that we had discovered from our EFA, and formally modelled the structure (a bifactor model, with one global factor, and two subsequent group factors) in SEM. We wanted to ascertain the goodness of fit, map the standardised loadings back to the brain, and then ask how these observed factors of brain change were associated with *APOE* status and cognitive declines. Factor loadings from the CFA are plotted onto the cortical surface in Figure 4 (and are also provided in Supplementary Table 13). The CFA showed good fit to the data (Supplementary Table 14), and the brain cortical ROIs’ slope loadings on the three slope factors were highly commensurate with the exploratory findings (Supplementary Figure 7). The global factor showed general loadings across the cortex, whereas Factor 1 pertained clearly to frontal and temporal regions and Factor 2 related to occipital and parietal cortex; we shall henceforth refer to these as fronto-temporal and posterior-parietal factors, respectively. These factors explained a total of 86% of the total variance in regional slopes (general factor = 63%, fronto-temporal factor = 16%, and occipito-parietal factor = 7%). It is noteworthy that, although we refer to this largest factor as ‘general’ (as it is indicated by all but one region - pericalcarine), the actual magnitude of the CFA-estimated loadings ranged from 0.348 to 1.00, though most were large (M = 0.79, SD = 0.18; Supplementary Table 13). Loadings were strongest (>0.80) in the insula, dorsolateral and medial prefrontal, cingulate, lateral temporal and inferior parietal areas. In contrast, ventrolateral frontal, medial temporal, superior parietal and cuneal cortex showed relatively lower loadings (range 0.348 to 0.670). Fronto-temporal factor and occipito-parietal factor — which are independent of (orthogonal to) the general factor—were negatively correlated (*r* = −0.254, *p* < 0.001), indicating that participants exhibiting greater fronto-temporal decline tended to show less occipito-parietal decline, for any given level of global cortical volume change. Females and males did not differ in their trajectories of any aspect of cortical change (general = 0.020, *p* = 0.726, fronto-temporal = 0.020, *p* = 0.735, occipito-parietal = −0.005, *p* = 0.911).

**Figure 4.**
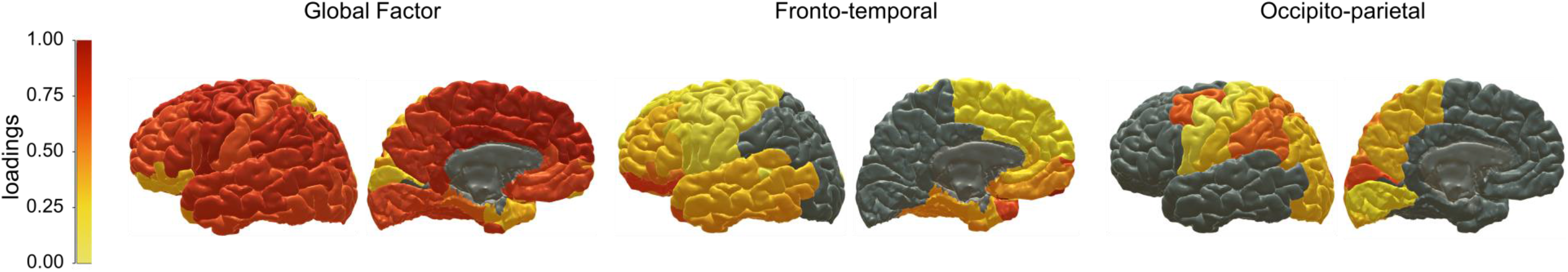
Cortical patterning of factors of cortical ageing *Note.* Warmer colours denote stronger standardised factor loadings of each ROI volume on each of the factors of cortical change estimated in the confirmatory factor analysis (see Supplementary Table 13). Grey colour denotes no loading. All ROIs except the pericalcarine cortex loaded onto the general factor, with subsequent loadings on Factors 1 (fronto-temporal) or Factor 2 (occipito-parietal) indicating that these ROIs exhibited common ageing trajectories in addition to the global pattern of overall cortical decline.

### Are patterns of cortical ageing related to *APOE* e4 status and cognitive decline?

An example of the multivariate SEM correlating cortical and cognitive ageing is shown in Figure 5. Measurement models of the cognitive domains, and the multivariate models correlating MRI with *APOE* and cognitive measures all showed good fit to the data (Supplementary Tables 14 and 15). *APOE* e4 carriers exhibited steeper atrophy for the general factor (*r* = −0.100, *p* = 0.038), but the statistical significance of this association did not survive FDR correction (Table 2). *APOE* e4 carriers did not show significantly steeper cortical atrophy on either fronto-temporal or occipito-parietal factors (*r* = −0.044, *p* = 0.414; *r* = 0.012, *p* = 0.774, respectively).

**Figure 5.**
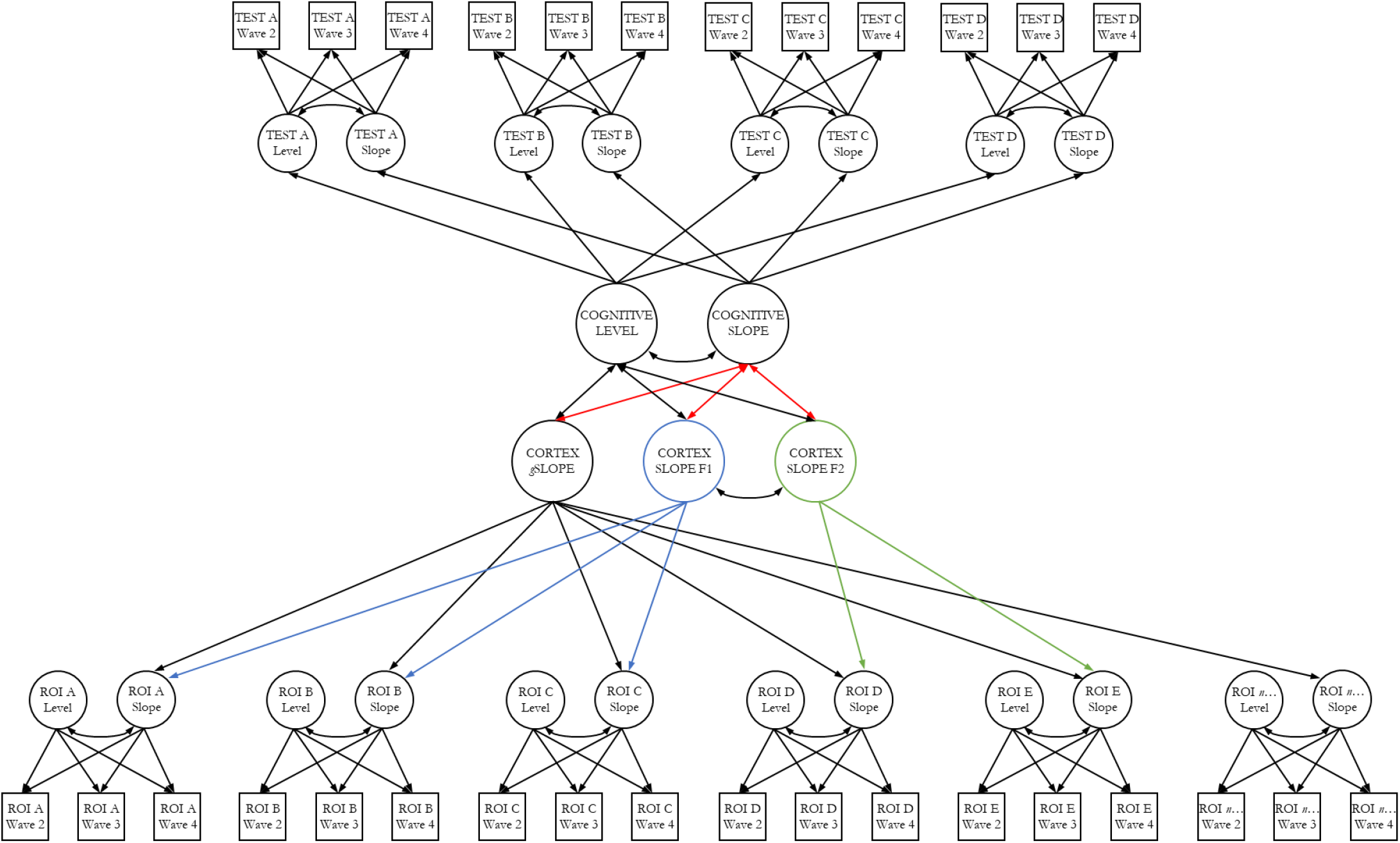
Modelling the coupled changes between cortical factors and cognitive domains. *Note.* An example of a multivariate latent growth curve model assessing associations between cortical and cognitive changes. The top half of the model illustrates how the intercept and slope of a given cognitive domain is indicated by the individual intercept and slope of multiple individual cognitive tests, tested on 3 occasions. The bottom half of the model illustrates how the 3 factors of cortical change are differentially indicated by the individual slopes of each of 34 cortical regions of interest (ROI). Residual correlations among manifest variables not shown. ROI intercept factors were freely estimated and allowed to correlate with all latent factors (not shown to reduce figure complexity). Red paths denote associations of interest, between cognitive and cortical changes. The two ‘secondary’ factors of cortical volume (F1 and f2) are orthogonal to the general slope of cortical volume, and negatively correlated with each other (*r* = −0.254, *p* < 0.001).

**Table 2.** Associations between factors of cortical change and *APOE* status, and changes in cognitive domains.

|  | Global | Fronto-temporal<br>Factor | Occipito-parietal<br>Factor |
| --- | --- | --- | --- |
| *APOE* e4 carrier | -0.100 (0.038) | -0.044 (0.414) | 0.012 (0.774) |
| *g* | **0.431 (<0.001)** | -0.024 (0.738) | -0.072 (0.212) |
| Visuospatial | **0.415 (0.002)** | -0.139 (0.286) | -0.143 (0.229) |
| Processing Speed | **0.369 (<0.001)** | -0.010 (0.893) | -0.066 (0.256) |
| Memory | **0.363 (<0.001)** | 0.066 (0.352) | -0.026 (0.644) |

*Note.* Standardised estimates (p-values) are reported for associations between the three factors of cortical volumetric change (global, fronto-temporal and posterior-parietal), *APOE* status (where 1 = at least 1 × e4 allele), and the three cognitive domains (visuospatial, processing speed and memory). Bold typeface denotes FDR- $q < 0.05$ .

Associations between the changes in cognitive domains and brain cortical factors are shown in Table 2. Trajectories of cognitive change in this sample have already been characterised and reported previously^28^. Greater cortical volumetric decline at the global level was significantly associated with declines in general cognitive ability (*g*) (*r* = 0.431, *p* < 0.001), visuospatial ability (*r* = 0.415, *p* = 0.002), processing speed (*r* = 0.383, *p* < 0.001), and memory ability (*r* = 0.372, *p* < 0.001). No other correlations among cognitive changes and either of the secondary factors of cortical change were FDR-significant (all *r*s ≤ |0.143|).

## Discussion

In this cohort of community-dwelling older adults, assessed three times over the course of their 8^th^ decade, we identified three axes of cortical change, along which general and anatomically-localised atrophy occurs. That is, we identified three dimensions of cortical atrophy which described clusters of areas exhibiting correlated rates of ageing. Together, these three patterns explained 83% of the individual differences in cortical ageing across 34 bilateral ROIs. Changes across the cortex were, in general, strongly correlated, and a general factor accounted for 63% of volumetric changes across the 34 ROIs. This general factor was associated with declines in a measure of general cognitive function (‘g’), which was more strongly driven by processing speed and memory than by visuospatial declines. Two additional, mildly-negatively-correlated factors existed on an anterior-posterior axis, and together explained an additional 20% of the variation in cortical ageing. Thus, for any given level of global cortical ageing, individuals tend additionally to experience either more fronto-temporal or more occipito-parietal cortical ageing. However, change across either of these additional dimensions of cortical atrophy were not significantly related to any latent measures of cognitive decline, beyond the principal axis of cortical atrophy.

Just as differential psychology has determined that a single psychological factor may largely underlie age-related declines across multiple cognitive tests^47^, so the current findings indicate that a large proportion of cortical atrophy occurs across a single dimension – and furthermore, that general cortical and general cognitive ageing are significantly coupled. By identifying that changes in both cognitive abilities and cortical volumes appear to occur along general – and correlated – statistical dimensions, these findings and similar approaches represent an important step in guiding ongoing research into the neuroanatomical correlates and potential underlying mechanisms of cognitive ageing. With respect to the interpretation of the general factor of cortical change, we note that fronto-temporal changes are more representative of overall cortical change: these lobes comprise a larger proportion of overall cortex, and our general factor was indicated by a far greater proportion of frontal and temporal parcels (k = 22 parcels) than parietal and occipital parcels (k = 12 parcels), and showed some subtle variation in loading magnitudes (though most were uniformly large). As such, we interpret the general cortical factor - and its correlations with cognitive ageing - as pertaining more strongly to fronto-temporal than occipito-parietal cortex. Thus, our findings do not mean that tests thought to tap our higher cognitive functioning are not related to frontal and temporal declines; rather, much of the cognitively-meaningful change in these brain regions is reflected in the first dimension of cortical ageing. Having additional – subtle - change anteriorly or posteriorly above this general dimension did not appear to account for significantly more cognitive change, though greater statistical power and a less healthy population may aid the reliable estimation of these secondary factors’ associations with cognitive ageing.

Strikingly, the patterns of structural covariance in cortical *changes* departed substantially from those observed in concurrent patterns of structural covariance among *levels* at baseline.

Correlations among baseline ROI volumes were moderate (average *r* = 0.350, SD = 0.178) in magnitude, yet markedly weaker than correlations among rates of longitudinal atrophy (average *r* = 0.805, SD = 0.252). There was only a subtle resemblance between the factors structure for level and change; a result that underscores previous claims for the theoretical and empirical strengths of using longitudinal data to inform accounts of ageing^5–7^. Only longitudinal data can be used to directly estimate the dimensionality of change over time, and to distinguish lifelong patterns of structural covariance from patterns that are specific to ageing.

We showed that the factor structure of cortical decline did not change when we removed those who had some indication of dementia, or stroke, suggesting that these patterns are also present in ostensibly ‘healthy’ ageing. The fronto-temporal pattern is partly consistent with frontal ageing accounts of cognitive decline^48,49^, and with the partial overlap between fronto-temporal ageing in healthy and Alzheimer’s-type patterns^50^. It is also notable that these three factors of change are similar to three of the five aspects of cortical change identified using a similar factor analytic method, in participants with MCI^18^. They found that individuals with greater frontal and temporal change, but not a more posterior pattern, were significantly more likely to convert to a clinical diagnosis of Alzheimer’s disease. However, the pattern of global cortical ageing is far more accentuated in our sample of healthy older adults – and the anterior and posterior patterns substantially weaker – suggesting a stark contrast between clinical patients and the sub-clinical majority of the ageing population. Nevertheless, these anterior and posterior patterns are similar to neurobiological patterns of dementia subtypes such as fronto-temporal and posterior cortical degeneration. Fronto-temporal lobar degeneration is a well-known pattern in pathological ageing, representing one of the main causes of dementia (accounting to up to 10% of all dementias^51^). In contrast, posterior cortical atrophy (PCA) is characterised by selective decline in functions mainly reliant on parietal and occipital brain regions^52,53^. It is estimated to account for around 8% of Alzheimer’s disease cases^54^, though its prevalence is currently unknown, and it is relatively under-researched^52^. These apparent similarities should be interpreted with additional caution though, given that none of the cortical changes were significantly steeper in carriers of the *APOE* e4 allele, which is a well-known risk factor for late-life dementia^55^.

The study has limitations. Whereas it is tempting to draw parallels between the patterns of cortical atrophy identified here and those observed in clinical subtypes, volumetric atrophic effects likely reflect numerous ongoing processes. On the other hand, we were unable to rule out the influence of nascent clinical neurodegeneration on the patterns of cortical atrophy discovered here, which we based on self-reported dementia and MMSE scores. It is possible that these additional profiles (anterior and posterior) reflect nascent and separable pathological neurodegenerative processes. Whereas we consider it unlikely that our results are predominantly driven by such effects (given the general prevalence of these cases in the population^51,54^), neurostructural hallmarks may be detectable prior to the onset of cognitive impairments^56^. It may therefore be of interest to test whether these additional patterns of cortical change, which we identified in the eighth decade of life, predict future cognitive trajectories as longitudinal testing continues into older ages. There are also many other aspects of brain structure and function that were not measured which may shed further light on any potential similarities and differences with pathological ageing. Moreover, the present results indicate that these additional patterns are relatively subtle among relatively healthy and range restricted^57^ group of older adults. As such, it is possible that these are underestimates of the true extent to which these patterns are present at the population level. Nevertheless, we note that our ability to account for all available data in our models ensured that we did not overly bias our results toward the most cognitively able participants who returned for all three neuroimaging visits, as we might have done modelling completers only. The impact that these two approaches have on model-implied trajectories has been neatly illustrated elsewhere, using cognitive data from the present cohort^58^.

In summary, our analyses have revealed that i) cortical ageing occurs partly as a coordinated mantle-wide process, and beyond that, as either greater fronto-temporal or greater occipito-parietal atrophy; ii) this same pattern is not readily apparent from cross-sectional data; iii) the major, single/general, axis of cortical atrophy (indicated by a greater number of fronto-temporal regions) is related to general cognitive decline, and processing speed and memory cognitive ability domains.

## Author Contributions

SRC, IJD and ETD designed the analysis. SRC conducted the analysis and drafted the work. MEB, JMW and IJD conceived the study design, SRC, MH, MVH, JC, AT and JMW collected or analysed the brain MRI and/or cognitive data. All authors critically revised the work and have approved the submitted version.

## Data and Code Availability

Materials, code, and associated protocols will be made available upon reasonable request to the corresponding author. The data analysed in this study is not publicly available as it contains data that could compromise participant consent and confidentiality, but can be requested via a data access request to the Lothian Birth Cohorts research group.

## Acknowledgements

We thank the Lothian Birth Cohort 1936 members who took part in this study, and Lothian Birth Cohort 1936 research team members who collected, entered and checked data used in this manuscript. The LBC1936 and this research are supported by Age UK (Disconnected Mind project) and by the UK Medical Research Council [MRC; G0701120, G1001245, MR/M013111/1, MR/R024065/1]. SRC, SJR, MEB, IJD and EMT-D were also supported by a National Institutes of Health (NIH) research grant R01AG054628. MH, AMM, HCW, JMW, IJD are also supported by a Wellcome Trust Strategic Award (Ref 104036/Z/14/Z). JMW receives funding from the UK Dementia Research Institute (funded by the MRC, Alzheimer’s Society and Alzheimer’s Research UK), the Fondation Leducq, the BHF Edinburgh Centre for Research Excellence and the Row Fogo Charitable Trust. MCVH is funded by the Row Fogo Charitable Trust (grant No. BROD.FID3668413). EMT-D is a member of the Population Research Center at the University of Texas at Austin, which is supported by NIH center grant P2CHD042849. TCR is a member of the Alzheimer Scotland Dementia Research Centre supported by Alzheimer Scotland.

