## Supplementary Material for "Three major dimensions of human brain cortical ageing in relation to cognitive decline across the 8^th^ decade of life"

**Supplementary Table 1.** Baseline (Wave 2) characteristics of participants who attended neuroimaging at Wave 2, Wave 3 and Wave 4.

|  | **Wave 2** |  | **Wave 3** |  | **Wave 4** |  |
| --- | --- | --- | --- | --- | --- | --- |
|  | **M (SD)** | **N** | **M(SD)** | **N** | **M(SD)** | **N** |
| **Age (years)** | 72.50 (0.70) | 731 | 72.50 (0.71) | 488 | 72.52 (0.72) | 386 |
| **Sex M:F** | 388:343 | 731 | 260:228 | 488 | 206:180 | 386 |
| ***APOE* e4 carriers (N:Y)** | 490:205 | 695 | 320:144 | 464 | 261:107 | 368 |
| **Cortical Volume (mm^3^)** | 398508 (37288) | 629 | 403474 (36824) | 453 | 403962 (37215) | 359 |
| **Matrix Reasoning** | 13.36 (4.92) | 729 | 13.63 (4.91) | 486 | 14.08 (4.84) | 384 |
| **Block Design** | 34.00 (10.07) | 729 | 34.74 (10.29) | 488 | 34.98 (10.24) | 385 |
| **Spatial Span Total** | 14.75 (2.77) | 728 | 14.83 (2.71) | 484 | 14.89 (10.24) | 385 |
| **Symbol Search** | 24.62 (6.19) | 729 | 25.09 (5.97) | 486 | 25.68 (5.47) | 383 |
| **Digit Symbol** | 56.29 (12.42) | 729 | 57.45 (12.08) | 486 | 58.54 (11.79) | 383 |
| **Inspection Time** | 111.37 (11.75) | 714 | 111.92 (11.76) | 478 | 112.69 (11.59) | 380 |
| **4-Choice Reaction Time** | 0.65 (0.09) | 730 | 0.64 (0.08) | 487 | 0.63 (0.08) | 385 |
| **Logical Memory** | 74.45 (17.93) | 729 | 75.47 (17.29) | 488 | 77.02 (16.56) | 386 |
| **Verbal Pairs** | 27.18 (9.57) | 713 | 27.52 (9.59) | 477 | 28.50 (9.19) | 379 |
| **Digit Span Backwards** | 7.86 (2.32) | 731 | 7.90 (2.31) | 488 | 8.06 (2.29) | 386 |

*Note.* Showing data **at Wave 2** for participants who attended the imaging assessment across Wave 2 to 4 (brain imaging data was not collected at Wave 1 of the study).

**Supplementary Table 2.** Baseline (Wave 2) characteristics of participants who attended Wave 2 but not Wave 3, and Wave 3 but not Wave 4.

|  | **MRI Wave 2, but not Wave 3** | | **MRI Wave 3, but not Wave 4** | |
| --- | --- | --- | --- | --- |
|  | **M (SD)** | **N** | **M(SD)** | **N** |
| **Age (years)** | 72.52 (0.71) | 250 | 72.49 (0.69) | 132 |
| **Sex M:F** | 130:120 | 250 | 68:64 | 132 |
| ***APOE* e4 carriers (N:Y)** | 175:63 | 236 | 81:44 | 125 |
| **Cortical Volume (mm^3^)** | 385726 (35485) ^a^ | 176 | 400607 (34853) | 116 |
| **Matrix Reasoning** | 12.72 (4.90) ^a^ | 250 | 12.27 (4.86)^b^ | 132 |
| **Block Design** | 32.37 (9.46) ^a^ | 248 | 33.47 (10.02) | 132 |
| **Spatial Span Total** | 14.57 (2.90) | 250 | 14.48 (2.53) | 130 |
| **Symbol Search** | 23.62 (6.59) ^a^ | 249 | 23.50 (6.83) ^b^ | 132 |
| **Digit Symbol** | 53.83 (12.82) ^a^ | 249 | 53.97 (12.19) ^b^ | 132 |
| **Inspection Time** | 110.35 (11.68) | 241 | 109.52 (12.21) ^b^ | 126 |
| **4-Choice Reaction Time** | 0.66 (0.10) ^a^ | 250 | 0.67 (0.09) ^b^ | 132 |
| **Logical Memory** | 71.83 (19.43) ^a^ | 248 | 71.20 (19.20) ^b^ | 132 |
| **Verbal Pairs** | 26.28 (9.60) | 241 | 24.84 (10.28) ^b^ | 128 |
| **Digit Span Backwards** | 7.73 (2.36) | 250 | 7.41 (2.28) ^b^ | 132 |

*Note.* Showing data for participants **at Wave 2** who attended neuroimaging at Wave 2 but not Wave 3, and Wave 3 but not Wave 4. ^a^ denotes a significant difference (*p* < 0.05, uncorrected) in the imaging sample between Wave 2 values for those who attended Wave 2 *and* 3 versus those who attended Wave 2 but not Wave 3 – that is, a difference between the values shown in Supplementary Table 1, column 2, and column 1 this table; ^b^ denotes a significant difference (*p* < 0.05, uncorrected) in the imaging sample between Wave 2 values for those who attended Wave 3 *and* 4 versus those who attended Wave 3 but not Wave 4 – that is, a difference between the values shown in Supplementary Table 1, column 3, and column 2 this table.

**Supplementary Table 3.** Estimates of regional cortical change.

|  | ***Hemi*** | ***β*** | **SE** | ***p*** | **% loss p.a.** |
| --- | --- | --- | --- | --- | --- |
| ***GLOBAL*** |  |  |  |  |  |
| Volume (cm^3^) |  | -3.475 | 0.130 | <0.001 | 0.872 |
| ***REGIONAL VOLUME*** |  |  |  |  |  |
| Banks STS | L | -1.580 | 0.123 | 0.000 | -0.761 |
|  | R | -1.649 | 0.105 | 0.000 | -0.840 |
| Caudal ACC | L | -0.306 | 0.109 | 0.005 | -0.211 |
|  | R | -0.705 | 0.123 | 0.000 | -0.410 |
| Caudal Middle Frontal | L | -4.505 | 0.294 | 0.000 | -0.844 |
|  | R | -3.889 | 0.272 | 0.000 | -0.772 |
| Cuneus | L | -2.260 | 0.137 | 0.000 | -0.864 |
|  | R | -2.828 | 0.135 | 0.000 | -1.026 |
| Entorhinal | L | -1.947 | 0.202 | 0.000 | -1.059 |
|  | R | -1.982 | 0.163 | 0.000 | -1.147 |
| Frontal Pole | L | -1.596 | 0.073 | 0.000 | -1.892 |
|  | R | -1.900 | 0.092 | 0.000 | -1.720 |
| Fusiform | L | -7.041 | 0.547 | 0.000 | -0.787 |
|  | R | -7.071 | 0.493 | 0.000 | -0.821 |
| Inferior Parietal | L | -13.632 | 0.543 | 0.000 | -1.215 |
|  | R | -15.137 | 0.605 | 0.000 | -1.141 |
| Inferior Temporal | L | -10.440 | 0.503 | 0.000 | -1.048 |
|  | R | -10.652 | 0.477 | 0.000 | -1.105 |
| Insula | L | -1.717 | 0.314 | 0.000 | -0.265 |
|  | R | -1.946 | 0.376 | 0.000 | -0.297 |
| Isthmus Cingulate | L | -1.506 | 0.145 | 0.000 | -0.604 |
|  | R | -1.419 | 0.131 | 0.000 | -0.620 |
| Lateral Occipital | L | -13.191 | 0.453 | 0.000 | -1.224 |
|  | R | -13.365 | 0.435 | 0.000 | -1.228 |
| Lateral Orbitofrontal | L | -6.154 | 0.370 | 0.000 | -0.908 |
|  | R | -5.546 | 0.365 | 0.000 | -0.837 |
| Lingual | L | -4.641 | 0.287 | 0.000 | -0.789 |
|  | R | -5.112 | 0.288 | 0.000 | -0.849 |
| Medial Orbitofrontal | L | -4.520 | 0.335 | 0.000 | -0.877 |
|  | R | -3.724 | 0.327 | 0.000 | -0.761 |
| Middle Temporal | L | -10.108 | 0.442 | 0.000 | -1.107 |
|  | R | -11.366 | 0.496 | 0.000 | -1.093 |
| Paracentral | L | -1.770 | 0.158 | 0.000 | -0.586 |
|  | R | -1.995 | 0.158 | 0.000 | -0.596 |
| Parahippocampal | L | -1.418 | 0.171 | 0.000 | -0.770 |
|  | R | -1.119 | 0.148 | 0.000 | -0.660 |
| IFG Pars Opercularis | L | -2.633 | 0.173 | 0.000 | -0.660 |
|  | R | -2.501 | 0.156 | 0.000 | -0.748 |
| IFG Pars Orbitalis | L | -2.402 | 0.127 | 0.000 | -1.246 |
|  | R | -3.007 | 0.133 | 0.000 | -1.301 |
| IFG Pars Triangularis | L | -2.654 | 0.151 | 0.000 | -0.897 |
|  | R | -3.153 | 0.162 | 0.000 | -0.901 |
| Pericalcarine | L | -1.027 | 0.082 | 0.000 | -0.594 |
|  | R | -1.027 | 0.098 | 0.000 | -0.531 |
| Postcentral | L | -5.640 | 0.389 | 0.000 | -0.659 |
|  | R | -5.378 | 0.398 | 0.000 | -0.667 |
| Posterior Cingulate | L | -1.390 | 0.184 | 0.000 | -0.502 |
|  | R | -1.610 | 0.169 | 0.000 | -0.587 |
| Precentral | L | -9.581 | 0.572 | 0.000 | -0.846 |
|  | R | -9.874 | 0.557 | 0.000 | -0.875 |
| Precuneus | L | -6.721 | 0.408 | 0.000 | -0.807 |
|  | R | -6.621 | 0.391 | 0.000 | -0.779 |
| Rostral ACC | L | -0.986 | 0.140 | 0.000 | -0.430 |
|  | R | -0.447 | 0.132 | 0.001 | -0.242 |
| Rostral Middle Frontal | L | -13.632 | 0.631 | 0.000 | -1.066 |
|  | R | -13.553 | 0.616 | 0.000 | -1.018 |
| Superior Frontal | L | -15.113 | 0.810 | 0.000 | -0.790 |
|  | R | -14.234 | 0.769 | 0.000 | -0.774 |
| Superior Parietal | L | -13.907 | 0.662 | 0.000 | -1.177 |
|  | R | -12.900 | 0.638 | 0.000 | -1.095 |
| Superior Temporal | L | -8.502 | 0.459 | 0.000 | -0.872 |
|  | R | -8.426 | 0.461 | 0.000 | -0.880 |
| Supramarginal | L | -8.396 | 0.419 | 0.000 | -0.887 |
|  | R | -7.141 | 0.391 | 0.000 | -0.792 |
| Temporal Pole | L | -3.913 | 0.266 | 0.000 | -1.444 |
|  | R | -3.323 | 0.245 | 0.000 | -1.337 |
| Transverse Temporal | L | -1.032 | 0.055 | 0.000 | -1.043 |
|  | R | -0.808 | 0.045 | 0.000 | -1.050 |

*Note.* Unstandardised estimates of annual change in cortical regional volume. ACC: anterior cingulate cortex; L: left; R: right.

**Supplementary Table 4.** Loadings of an exploratory analysis using Schmid-Leiman transform of left hemisphere ROI growth curves.

| **Region of Interest** | ***g*** | **F1** | **F2** |
| --- | --- | --- | --- |
| Lateralorbitofrontal | 0.60 | 0.77 |  |
| Temporalpole | 0.65 | 0.75 |  |
| Entorhinal | 0.68 | 0.70 |  |
| Parsorbitalis | 0.72 | 0.69 |  |
| Frontalpole | 0.68 | 0.67 |  |
| Parstriangularis | 0.78 | 0.59 |  |
| Insula | 0.77 | 0.58 |  |
| Rostralmiddlefrontal | 0.81 | 0.56 |  |
| Parahippocampal | 0.80 | 0.53 |  |
| Rostralanteriorcingulate | 0.83 | 0.51 |  |
| Superiortemporal | 0.83 | 0.50 |  |
| Middletemporal | 0.83 | 0.49 |  |
| Fusiform | 0.83 | 0.49 |  |
| Parsopercularis | 0.83 | 0.48 |  |
| Inferiortemporal | 0.83 | 0.47 |  |
| Medialorbitofrontal | 0.84 | 0.45 | 0.31 |
| Superiorfrontal | 0.84 | 0.41 | 0.35 |
| Transversetemporal | 0.84 | 0.38 | 0.38 |
| Caudalanteriorcingulate | 0.83 | 0.38 | 0.38 |
| Caudalmiddlefrontal | 0.84 | 0.36 | 0.41 |
| Precentral | 0.84 | 0.36 | 0.40 |
| Cuneus | 0.58 |  | 0.78 |
| Superiorparietal | 0.69 |  | 0.71 |
| Lateraloccipital | 0.74 |  | 0.68 |
| Precuneus | 0.75 |  | 0.66 |
| Lingual | 0.75 |  | 0.64 |
| Inferiorparietal | 0.77 |  | 0.64 |
| Paracentral | 0.77 |  | 0.62 |
| Isthmuscingulate | 0.79 |  | 0.59 |
| Posteriorcingulate | 0.82 |  | 0.53 |
| Postcentral | 0.82 |  | 0.52 |
| Supramarginal | 0.83 |  | 0.48 |
| Banksttsts | 0.83 |  | 0.47 |
| Pericalcarine |  |  |  |

*Note.* Loadings for left hemisphere ROIs; values <0.3 not shown (pericalcarine does not load on any of the three factors). Eigenvalues are: *g* = 20.1, F1 = 6.60, F2 = 6.00. Prior to the S-L transformation, general factor loadings of F1 and F2 on *g* were constrained to equality, and F1 ~ F2 *r* = 0.544. The S-L transform takes a factor or PC solution, transforms it to an oblique solution, factors the oblique solution to find a higher order (*g*) factor, and then residualizes *g* out of the group factors.

**Supplementary Table 5.** Loadings of an exploratory analysis using a three-factor Schmid-Leiman transform of left hemisphere ROI growth curves.

| **Region of Interest** | ***g*** | **F1** | **F2** | **F3** |
| --- | --- | --- | --- | --- |
| Lateralorbitofrontal | 0.96 |  |  |  |
| Temporalpole | 0.99 |  |  |  |
| Entorhinal | 0.95 |  |  |  |
| Parsorbitalis | 1 |  |  |  |
| Frontalpole | 0.97 |  |  |  |
| Parstriangularis | 0.98 |  |  |  |
| Insula | 0.98 |  |  |  |
| Rostralmiddlefrontal | 0.97 |  |  |  |
| Parahippocampal | 0.93 |  |  |  |
| Rostralanteriorcingulate | 0.94 |  | 0.33 |  |
| Superiortemporal | 0.94 |  | 0.34 |  |
| Middletemporal | 0.93 |  | 0.35 |  |
| Fusiform | 0.92 |  | 0.36 |  |
| Parsopercularis | 0.93 |  | 0.36 |  |
| Inferiortemporal | 0.92 |  | 0.39 |  |
| Medialorbitofrontal | 0.91 |  | 0.41 |  |
| Superiorfrontal | 0.89 |  | 0.46 |  |
| Transversetemporal | 0.87 |  | 0.49 |  |
| Caudalanteriorcingulate | 0.84 |  | 0.52 |  |
| Caudalmiddlefrontal | 0.85 |  | 0.52 |  |
| Precentral | 0.86 |  | 0.51 |  |
| Cuneus |  |  | 0.96 |  |
| Superiorparietal | 0.44 |  | 0.89 |  |
| Lateraloccipital | 0.51 |  | 0.85 |  |
| Precuneus | 0.55 |  | 0.84 |  |
| Lingual | 0.55 |  | 0.83 |  |
| Inferiorparietal | 0.59 |  | 0.81 |  |
| Paracentral | 0.61 |  | 0.77 |  |
| Isthmuscingulate | 0.63 |  | 0.77 |  |
| Posteriorcingulate | 0.73 |  | 0.68 |  |
| Postcentral | 0.75 |  | 0.66 |  |
| Supramarginal | 0.78 |  | 0.62 |  |
| Banksttsts | 0.78 |  | 0.62 |  |
| Pericalcarine | 0.96 |  |  |  |

*Note.* Loadings for left hemisphere ROIs; values <0.3 not shown. Eigenvalues are: *g* = 22.7, F1 = 0.0, F2 = 9.8, F3 = 1.3.

**Supplementary Table 6.** Loadings of an exploratory analysis using a four-factor Schmid-Leiman transform of left hemisphere ROI growth curves.

| **Region of Interest** | ***g*** | **F1** | **F2** | **F3** | **F4** |
| --- | --- | --- | --- | --- | --- |
| Lateralorbitofrontal | 0.95 |  |  |  |  |
| Temporalpole | 0.98 |  |  |  |  |
| Entorhinal | 0.98 |  |  |  |  |
| Parsorbitalis | 0.97 |  |  |  |  |
| Frontalpole | 0.91 | 0.40 |  |  |  |
| Parstriangularis | 0.92 |  |  |  |  |
| Insula | 0.94 |  |  |  |  |
| Rostralmiddlefrontal | 0.91 |  | 0.35 |  |  |
| Parahippocampal | 0.89 |  | 0.39 |  |  |
| Rostralanteriorcingulate | 0.88 |  | 0.44 |  |  |
| Superiortemporal | 0.88 |  | 0.45 |  |  |
| Middletemporal | 0.87 |  | 0.46 |  |  |
| Fusiform | 0.87 |  | 0.47 |  |  |
| Parsopercularis | 0.87 |  | 0.47 |  |  |
| Inferiortemporal | 0.86 |  | 0.49 |  |  |
| Medialorbitofrontal | 0.84 |  | 0.52 |  |  |
| Superiorfrontal | 0.82 |  | 0.55 |  |  |
| Transversetemporal | 0.80 |  | 0.59 |  |  |
| Caudalanteriorcingulate | 0.78 |  | 0.62 |  |  |
| Caudalmiddlefrontal | 0.77 |  | 0.62 |  |  |
| Precentral | 0.78 |  | 0.60 |  |  |
| Cuneus |  |  | 0.97 |  |  |
| Superiorparietal | 0.34 |  | 0.93 |  |  |
| Lateraloccipital | 0.42 |  | 0.90 |  |  |
| Precuneus | 0.46 |  | 0.89 |  |  |
| Lingual | 0.47 |  | 0.89 |  |  |
| Inferiorparietal | 0.50 |  | 0.86 |  |  |
| Paracentral | 0.52 |  | 0.82 |  |  |
| Isthmuscingulate | 0.56 |  | 0.84 |  |  |
| Posteriorcingulate | 0.65 |  | 0.76 |  |  |
| Postcentral | 0.66 |  | 0.73 |  |  |
| Supramarginal | 0.70 |  | 0.71 |  |  |
| Banksttsts | 0.71 |  | 0.70 |  |  |
| Pericalcarine |  |  |  | 1 |  |

*Note.* Loadings for left hemisphere ROIs; values <0.3 not shown. Eigenvalues are: *g* = 19.75, F1 = 0.62, F2 = 12.21, F3 = 1.27, F4 = 0.07.

**Supplementary Table 7.** Loadings of an exploratory analysis using Schmid-Leiman transform of right hemisphere ROI growth curves.

| **Region of Interest** | ***g*** | **F1** | **F2** |
| --- | --- | --- | --- |
| Lateralorbitofrontal | 0.47 | 0.84 |  |
| Temporalpole | 0.37 | 0.85 |  |
| Entorhinal | 0.48 | 0.84 |  |
| Parsorbitalis | 0.60 | 0.80 |  |
| Frontalpole | 0.34 | 0.65 |  |
| Parstriangularis | 0.65 | 0.75 |  |
| Insula | 0.55 | 0.76 |  |
| Rostralmiddlefrontal | 0.63 | 0.76 |  |
| Parahippocampal | 0.56 | 0.77 |  |
| Rostralanteriorcingulate | 0.58 | 0.81 |  |
| Superiortemporal | 0.63 | 0.78 |  |
| Middletemporal | 0.61 | 0.79 |  |
| Fusiform | 0.64 | 0.77 |  |
| Parsopercularis | 0.66 | 0.74 |  |
| Inferiortemporal | 0.63 | 0.78 |  |
| Medialorbitofrontal | 0.60 | 0.80 |  |
| Superiorfrontal | 0.68 | 0.72 |  |
| Transversetemporal | 0.69 | 0.69 |  |
| Caudalanteriorcingulate | 0.62 | 0.77 |  |
| Caudalmiddlefrontal | 0.70 | 0.68 |  |
| Precentral | 0.70 | 0.68 |  |
| Cuneus | 0.67 | 0.71 |  |
| Superiorparietal | 0.71 | 0.65 |  |
| Lateraloccipital | 0.73 | 0.59 | 0.34 |
| Precuneus | 0.70 | 0.67 |  |
| Lingual | 0.64 | 0.54 |  |
| Inferiorparietal | 0.73 | 0.59 | 0.34 |
| Paracentral | 0.74 | 0.56 | 0.38 |
| Isthmuscingulate | 0.71 | 0.53 | 0.38 |
| Posteriorcingulate | 0.74 | 0.46 | 0.49 |
| Postcentral | 0.74 | 0.43 | 0.51 |
| Supramarginal | 0.73 | 0.33 | 0.60 |
| Banksttsts | 0.71 |  | 0.68 |
| Pericalcarine |  |  | 0.59 |

*Note.* Loadings for right hemisphere ROIs; values <0.3 not shown (pericalcarine does not load on any of the three factors). Eigenvalues are: *g* = 20.1, F1 = 6.60, F2 = 6.00. Prior to the S-L transformation, general factor loadings of F1 and F2 on *g* were constrained to equality, and F1 ~ F2 *r* = 0.383. The S-L transform takes a factor or PC solution, transforms it to an oblique solution, factors the oblique solution to find a higher order (*g*) factor, and then residualizes *g* out of the group factors.

**Supplementary Table 8.** Comparisons of factor structure between left and right hemispheres as identified by the exploratory Schmid-Leiman analysis of ROI slopes and intercepts.

|  | ***LOADINGS*** | | |
| --- | --- | --- | --- |
| Correspondence | **General Factor** | **Factor 1** | **Factor 2** |
| ***SLOPES***  ***Left vs Right Hemisphere*** |  |  |  |
| **Pearson’s** | 0.78 | 0.70 | 0.61 |
| **Factor Congruence** | 0.99 | 0.87 | 0.73 |
| ***INTERCEPTS***  ***Left vs Right Hemisphere*** |  |  |  |
| **Pearson’s *r*** | 0.97 | 0.76 | 0.90 |
| **Factor Congruence** | 1.00 | 0.92 | 0.91 |
| ***SLOPES versus INTERCEPTS***  ***Left vs Left & Right vs Right Hemisphere*** | |  |  |
| **Pearson’s *r*** | L (0.05) R (0.04) | L (0.29) R (0.51) | L (0.32) R (0.67) |
| **Factor Congruence** | L (0.91) R (0.93) | L (0.74) R (0.86) | L (0.45) R (0.70) |

*Note.* Standardised loadings are reported; L = left hemisphere; R = right hemisphere.

**Supplementary Table 9.** Loadings of an exploratory factor analysis using Schmid-Leiman transform of left-right averaged ROI growth curves.

| **Region of Interest** | ***g*** | **F1** | **F2** |
| --- | --- | --- | --- |
| Lateralorbitofrontal | 0.53 | 0.80 |  |
| Temporalpole | 0.56 | 0.79 |  |
| Entorhinal | 0.60 | 0.77 |  |
| Frontalpole | 0.50 | 0.75 |  |
| Parsorbitalis | 0.68 | 0.73 |  |
| Rostralanteriorcingulate | 0.75 | 0.65 |  |
| Rostralmiddlefrontal | 0.75 | 0.64 |  |
| Parahippocampal | 0.75 | 0.64 |  |
| Medialorbitofrontal | 0.76 | 0.64 |  |
| Fusiform | 0.76 | 0.63 |  |
| Superiortemporal | 0.77 | 0.61 |  |
| Parstriangularis | 0.77 | 0.61 |  |
| Inferiortemporal | 0.78 | 0.60 |  |
| Middletemporal | 0.78 | 0.60 |  |
| Caudalanteriorcingulate | 0.78 | 0.60 |  |
| Insula | 0.78 | 0.59 |  |
| Parsopercularis | 0.79 | 0.57 |  |
| Superiorfrontal | 0.79 | 0.56 |  |
| Transversetemporal | 0.81 | 0.51 |  |
| Precentral | 0.81 | 0.50 | 0.31 |
| Caudalmiddlefrontal | 0.81 | 0.49 | 0.31 |
| Bankssts | 0.81 | 0.48 | 0.32 |
| Posteriorcingulate | 0.81 | 0.45 | 0.37 |
| Supramarginal | 0.82 | 0.43 | 0.38 |
| Postcentral | 0.82 | 0.40 | 0.41 |
| Isthmuscingulate | 0.78 | 0.31 | 0.47 |
| Paracentral | 0.81 | 0.31 | 0.50 |
| Inferiorparietal | 0.80 |  | 0.53 |
| Lingual | 0.78 |  | 0.55 |
| Precuneus | 0.78 |  | 0.59 |
| Lateraloccipital | 0.77 |  | 0.62 |
| Superiorparietal | 0.74 |  | 0.67 |
| Cuneus | 0.61 |  | 0.78 |
| Pericalcarine |  |  | 0.38 |

*Note.* Loadings for ROI volumes (average of left and right); values <0.3 not shown. Prior to the S-L transformation, general factor loadings of F1 and F2 on *g* were constrained to equality, and F1 ~ F2 *r* = 0.501. The S-L transform takes a factor or PC solution, transforms it to an oblique solution, factors the oblique solution to find a higher order (*g*) factor, and then residualizes *g* out of the group factors.

**Supplementary Table 10.** Comparisons of factor structure of ROI slopes and intercepts (left and right averaged volumes) identified by the exploratory Schmid-Leiman analysis.

|  | ***LOADINGS*** | | |
| --- | --- | --- | --- |
| Correspondence | **General Factor** | **Factor 1** | **Factor 2** |
| ***SLOPES versus INTERCEPTS*** | |  |  |
| **Pearson’s *r*** | 0.30 | 0.53 | 0.54 |
| **Factor Congruence** | 0.95 | 0.88 | 0.60 |

**Supplementary Table 11**. Comparison of Schmid-Leiman factor loadings of bilateral cortical volumes in the main sample and the dementia/MMSE sensitivity sample.

| **Correspondence** | **General Factor** | **Factor 1** | **Factor 2** |
| --- | --- | --- | --- |
| **SLOPES vs SLOPES** |  |  |  |
| **Pearson’s *r*** | >0.99 | 0.98 | 0.96 |
| **Factor Congruence** | 1.00 | 1.00 | 1.00 |
| **INTERCEPTS versus INTERCEPTS** | |  |  |
| **Pearson’s *r*** | 0.92 | 0.71 | 0.92 |
| **Factor Congruence** | 1.00 | 1.00 | 1.00 |

*Note.* Sensitivity sample removed individuals with a self-reported diagnosis of dementia at any wave or a score of MMSE <24. See also Supplementary Figure 5.

**Supplementary Table 12**. Comparison of Schmid-Leiman factor loadings of bilateral cortical volumes in the main sample and the stroke sensitivity sample.

| **Correspondence** | **General Factor** | **Factor 1** | **Factor 2** |
| --- | --- | --- | --- |
| **SLOPES vs SLOPES** |  |  |  |
| **Pearson’s *r*** | 0.91 | 0.82 | 0.77 |
| **Factor Congruence** | 1.00 | 0.92 | 0.84 |
| **INTERCEPTS versus INTERCEPTS** | |  |  |
| **Pearson’s *r*** | 0.95 | 0.90 | >0.99 |
| **Factor Congruence** | 1.00 | 1.00 | 1.00 |

*Note.* Sensitivity sample removed individuals with stroke at any wave. See also Supplementary Figure 6. Unique participants in the whole sample with a stroke at any wave was N = 119, this equated to a removal of N = 108, N = 64 and N = 48 participants with QC’d FreeSurfer data at Waves 2, 3 & 4, respectively (i.e. many with stroke attended >1 wave).

**Supplementary Table 13.** Loadings from a confirmatory factor analysis of ROI growth curves.

| **Region of Interest** | ***g*** | **F1** | **F2** |
| --- | --- | --- | --- |
| Lateralorbitofrontal | 0.39 | 0.92 |  |
| Temporalpole | 0.52 | 0.81 |  |
| Frontalpole | 0.65 | 0.76 |  |
| Parsorbitalis | 0.48 | 0.70 |  |
| Medialorbitofrontal | 0.82 | 0.57 |  |
| Fusiform | 0.83 | 0.56 |  |
| Parahippocampal | 0.67 | 0.54 |  |
| Rostralmiddlefrontal | 0.81 | 0.51 |  |
| Inferiortemporal | 0.78 | 0.51 |  |
| Middletemporal | 0.89 | 0.48 |  |
| Superiortemporal | 0.88 | 0.47 |  |
| Entorhinal | 0.45 | 0.46 |  |
| Parstriangularis | 0.90 | 0.45 |  |
| Rostralanteriorcingulate | 0.79 | 0.42 |  |
| Insula | 0.92 | 0.39 |  |
| Superiorfrontal | 0.97 | 0.26 |  |
| Transversetemporal | 0.97 | 0.25 |  |
| Caudalanteriorcingulate | 0.98 | 0.22 |  |
| Parsopercularis | 0.84 | 0.22 |  |
| Caudalmiddlefrontal | 0.97 | 0.17 | 0.08 |
| Postcentral | 0.76 | 0.15 | 0.31 |
| Precentral | 0.96 | 0.14 | 0.14 |
| Bankssts | 0.79 | 0.13 |  |
| Posteriorcingulate | 1.00 |  |  |
| Supramarginal | 0.94 |  | 0.08 |
| Paracentral | 0.92 |  | 0.39 |
| Isthmuscingulate | 0.91 |  |  |
| Inferiorparietal | 0.91 |  | 0.42 |
| Precuneus | 0.87 |  | 0.44 |
| Lingual | 0.80 |  | 0.26 |
| Lateraloccipital | 0.77 |  | 0.42 |
| Superiorparietal | 0.52 |  | 0.72 |
| Cuneus | 0.35 |  | 0.89 |
| Pericalcarine |  |  |  |

*Note.* Standardised factor loadings reported. Confirmatory factor analysis imposed the factor loading pattern identified from the exploratory analysis, but freely estimated loading magnitudes. Loadings of the paracentral, posterior cingulate and isthmus cingulate were initially found to load on the Schmid-Leiman Factor 1, but were non-significant in the CFA, and so were set to zero. F1 and F2 are orthogonal to the general factor and correlated with one another at *r* = -0.254 (*p* < 0.001).

**Supplementary Table 14***.* Model fit indices for CFAs for cognitive and cortical growth curves (separate measurement models).

|  | **RMSEA** | **CFI** | **TLI** | **SRMR** |
| --- | --- | --- | --- | --- |
| Cortical ROIs | 0.045 | 0.936 | 0.921 | 0.029 |
| *g* | 0.035 | 0.966 | 0.965 | 0.065 |
| Visuospatial | 0.029 | 0.933 | 0.992 | 0.037 |
| Processing Speed | 0.034 | 0.924 | 0.910 | 0.054 |
| Memory | 0.037 | 0.934 | 0.921 | 0.050 |

*Note.* RMSEA: root mean squared error of approximation, CFI: comparative fit index, TLI: Tucker Lewis Index, SRMR: standardized root mean square residual.

**Supplementary Table 15***.* Model fit indices for bifactor growth curve SEMs between cortical volumes, *APOE,* and cognitive functions.

|  | **RMSEA** | **CFI** | **TLI** | **SRMR** |
| --- | --- | --- | --- | --- |
| *APOE* status | 0.035 | 0.937 | 0.923 | 0.029 |
| *g* | 0.031 | 0.931 | 0.922 | 0.049 |
| Visuospatial | 0.037 | 0.933 | 0.919 | 0.048 |
| Processing Speed | 0.034 | 0.923 | 0.909 | 0.054 |
| Memory | 0.037 | 0.933 | 0.920 | 0.050 |

*Note.* RMSEA: root mean squared error of approximation, CFI: comparative fit index, TLI: Tucker Lewis Index, SRMR: standardized root mean square residual.

**Supplementary Figure 1.** Cortical regions of interest.
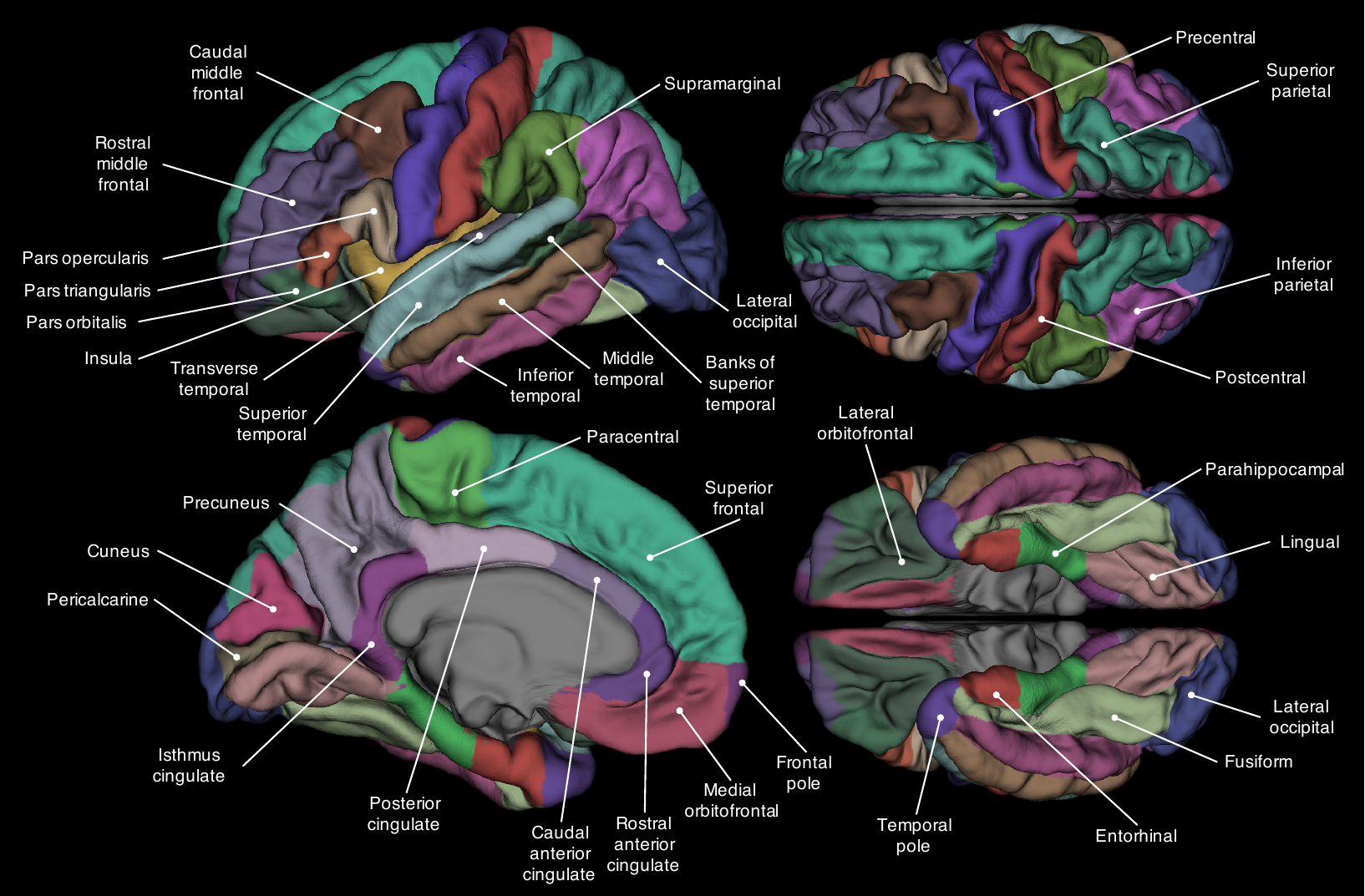

*Note.* Volumetric parcellation scheme used to identify regions of interest, according to the Desikan-Killiany atlas (Desikan et al., 2006).

**Supplementary Figure 2**.

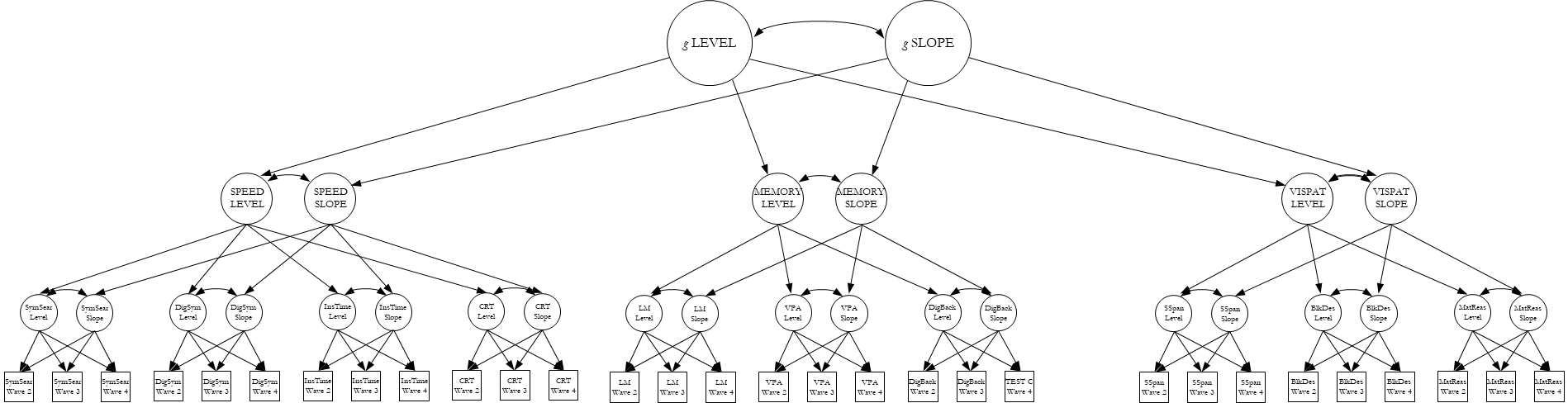

*Note.* Representation of a latent growth curve analysis in a hierarchical model of cognitive functioning. Manifest variables (i.e. measured test scores) are shown as squares, latent variables are shown as ellipses. The levels and slopes of the cognitive domains of processing speed (SPEED), memory and visuospatial ability (VISPAT) are indicated by the latent levels and changes of measured cognitive test scores across time. A superordinate latent measure of general cognitive ability (‘*g*’) accounts for the fact that the levels and changes across cognitive test scores in different domains are all positively correlated.

**Supplementary Figure 3**. Changes in regional cortical volume across the 8^th^ decade, by hemisphere.

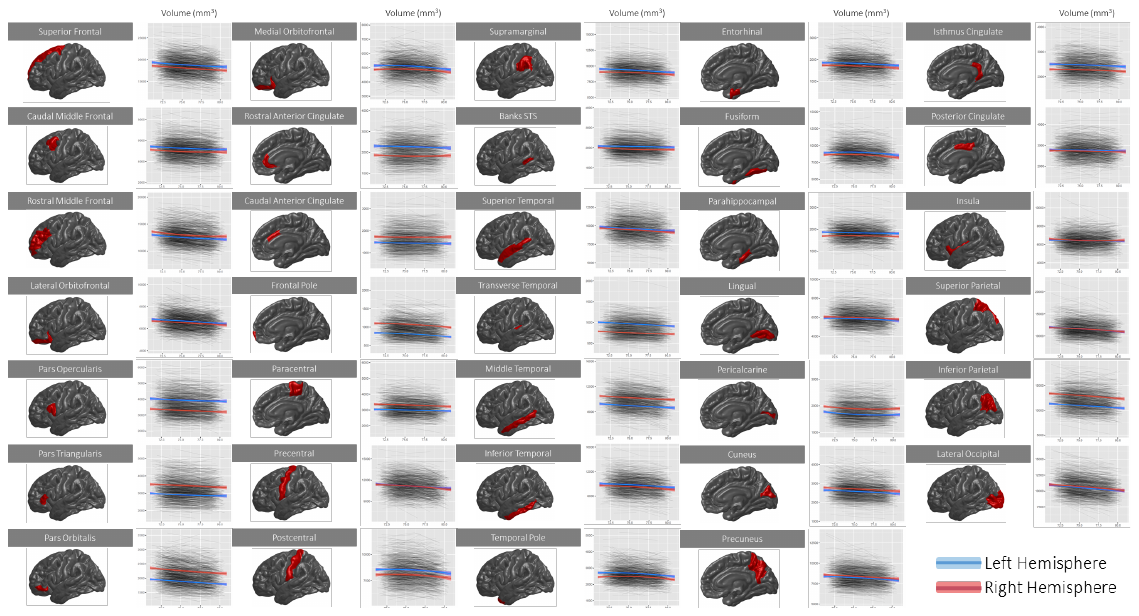

*Note.* Regional volumes (y-axis; mm^3^) are shown as age trajectories (x-axis; years) for each participant at each region of interest. Regression lines allowing a quadratic term, with 95% confidence intervals are shown for left and right hemispheres (blue and red, respectively). Unstandardised coefficients are reported in Supplementary Table 3.

**Supplementary Figure 4**. Correlation matrices and density plots of freely-estimated regional intercepts and slopes; left and right hemisphere comparison.

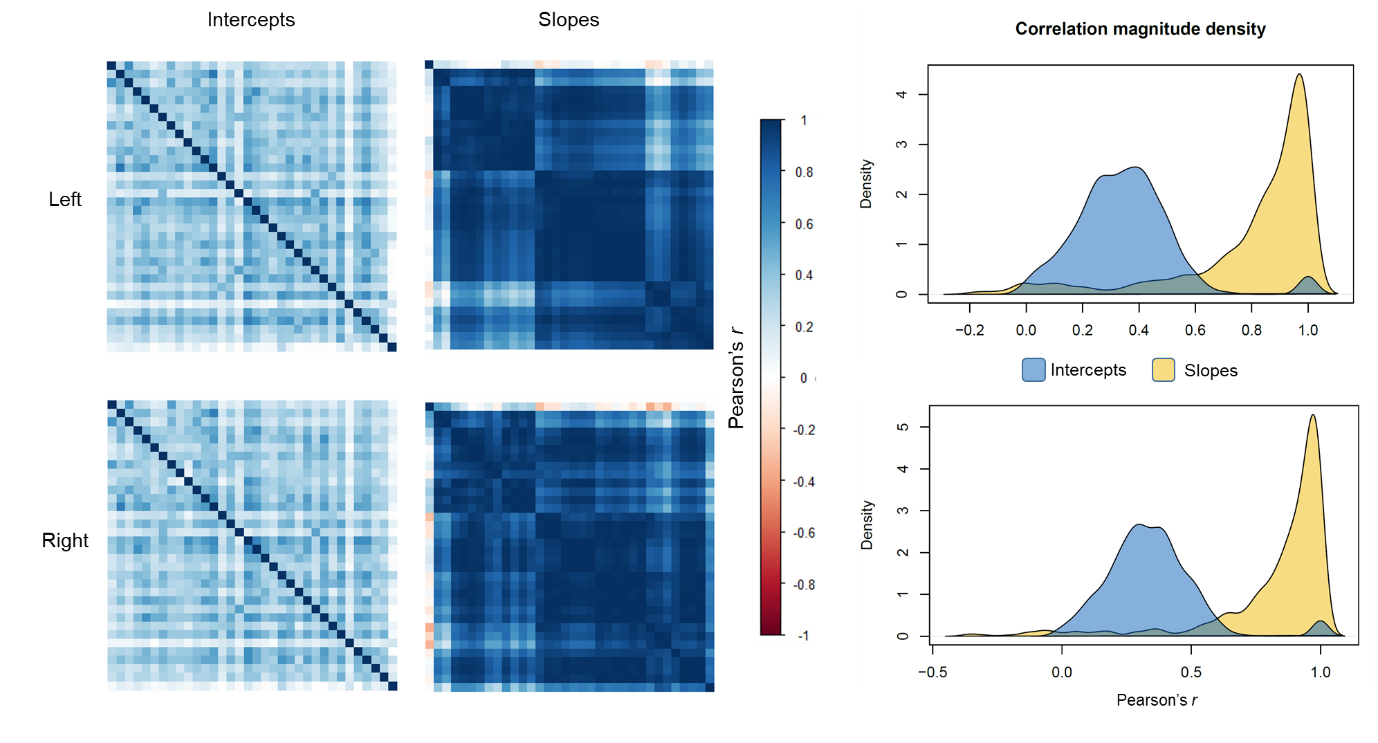

*Note.* **Left:** heatmaps of the correlations among freely-estimated latent intercepts and slopes for the left (top row) and right (bottom row) hemispheres; intercept axes are fixed according to the hierarchically-clustered slope matrix. **Right:** density plots of the correlation magnitudes among freely-estimated intercepts and slopes for the left (top) and right (bottom) hemispheres.

**Supplementary Figure 5.** Sensitivity analysis: difference in correlation magnitudes among intercepts and slopes of bilateral regional volumes before and after removing those with dementia or MMSE<24.

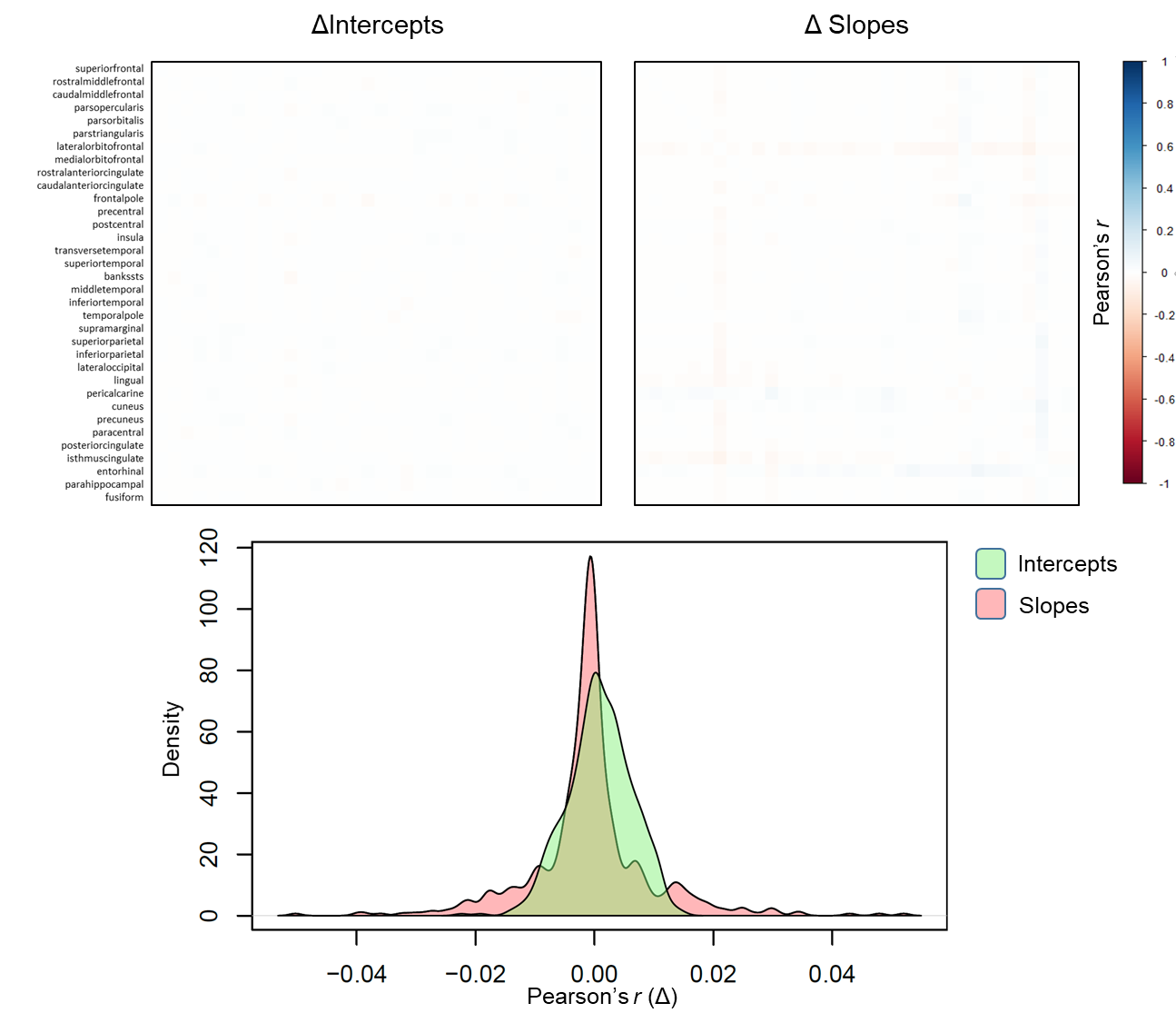

*Note.* Raw differences (Δ) between correlation magnitudes are shown for intercepts and slopes (top). The density plot shows the Δ distributions. Alongside the similarity of the resultant factor structure (see Supplementary Table 11), these results indicate that possible dementia cases are unlikely to have been driving the observed ageing patterns.

**Supplementary Figure 6.** Sensitivity analysis: difference in correlation magnitudes among intercepts and slopes of bilateral regional volumes before and after removing those with stroke.

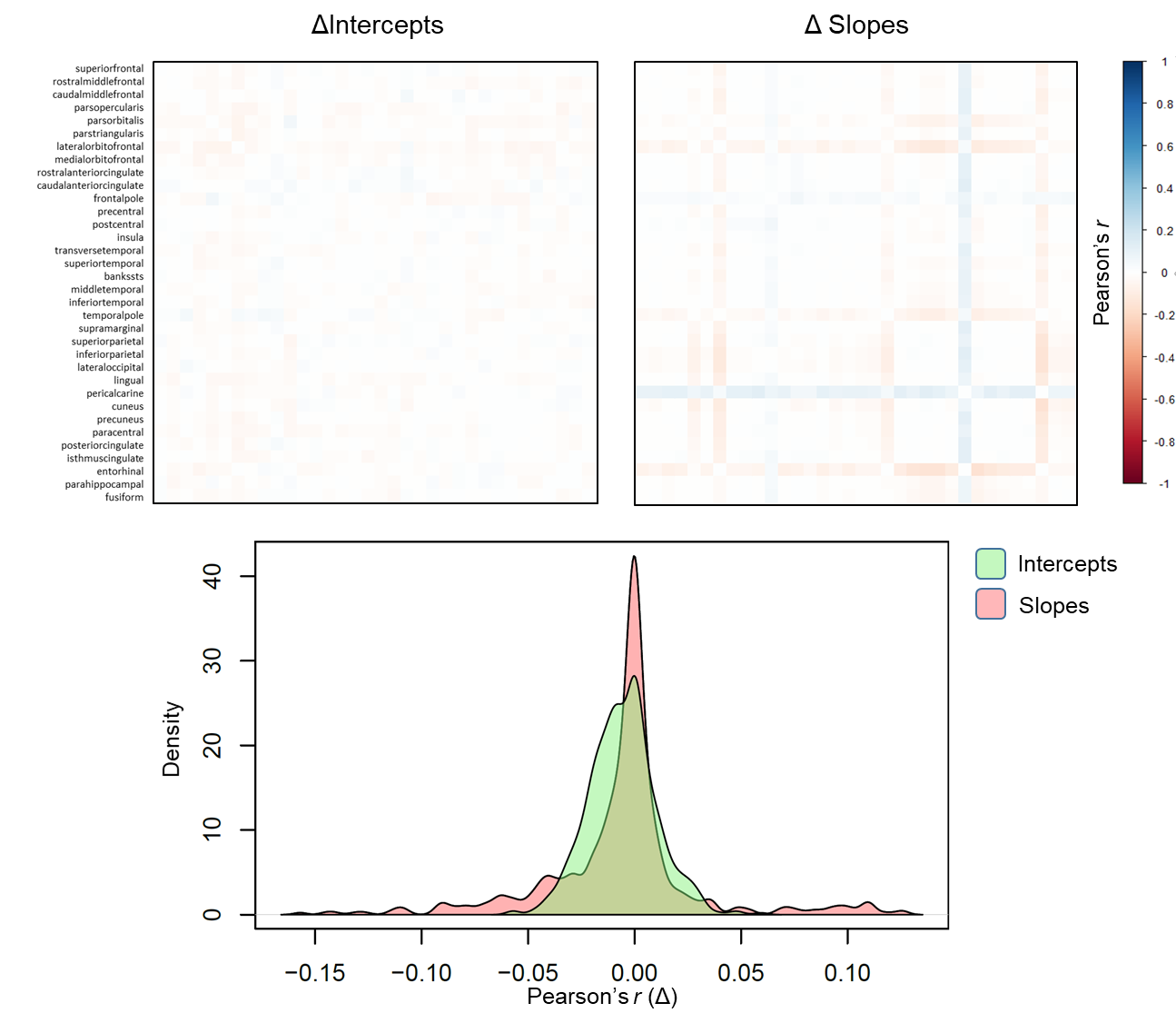

*Note.* Raw differences (Δ) between correlation magnitudes are shown for intercepts and slopes (top). The density plot shows the Δ distributions. Alongside the similarity of the resultant factor structure (see Supplementary Table 12), these results indicate that stroke cases are unlikely to have been driving the observed ageing patterns.

**Supplementary Figure 7***.* Correspondence between standardised loadings for factors of cortical change estimated via exploratory and confirmatory factor analyses.

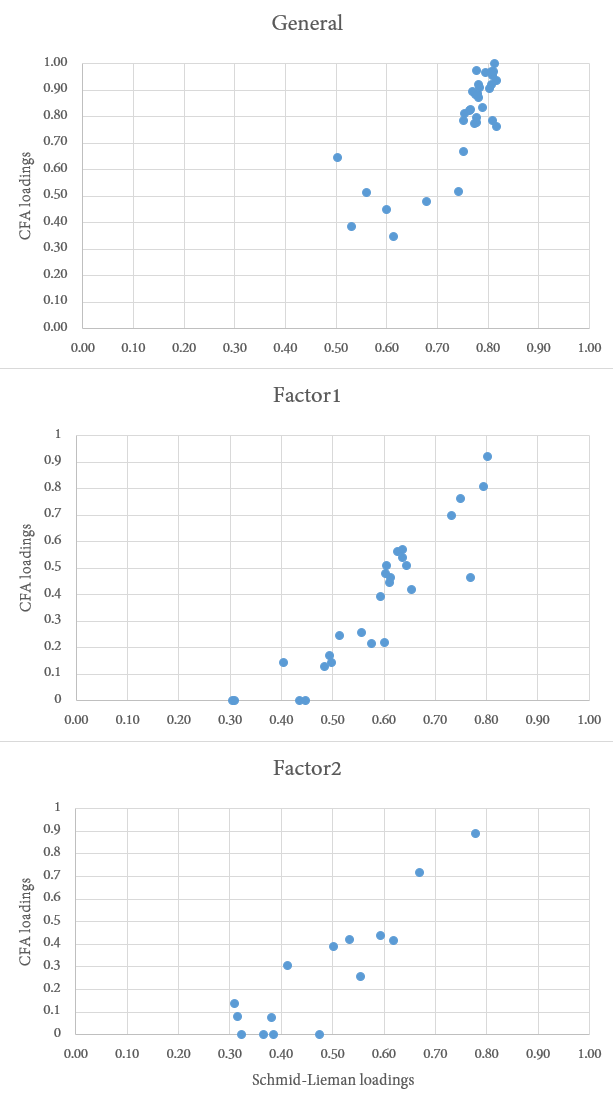

*Note.* X and Y axes denote standardised factor loadings from the exploratory Schmid-Leiman (X) and CFA (Y). Correlations (Pearson’s *r*) between EFA and CFA loadings for each factor were: General = 0.856, Factor 1 = 0.831, Factor 2 = 0.826.
